# Multimodal analysis of molecular remodeling in aging spleen identified global and cell type specific changes

**DOI:** 10.64898/2026.04.17.719305

**Authors:** Katarina Vlajic, Alison Luciano, Gennifer E Merrihew, Carla R. Sanchez, Michael Riffle, Kristine A. Tsantilas, Sahar Attar, Brian J. Beliveau, Mariya T. Sweetwyne, Michael J MacCoss, Gary Churchill, Devin K Schweppe

## Abstract

Aging reshapes the cellular and molecular landscape of mammalian tissues. These changes can be progressive, preceding linearly with age, or occur as abrupt transitions of the course of lifespan. To investigate the age-dependent cellular and molecular shifts we profiled matched proteomes and transcriptomes from male and female murine spleens across eight time points, from stable adults through late life. The spleen was chosen to integrate understanding of age-dependent changes associated with immune surveillance, inflammaging, and immune-related proteostasis. Male and female mice follow distinct aging trajectories particularly in protein–RNA correlation in late life, reflecting both compositional shifts and failure of post-transcriptional buffering. To investigate whether these changes could be attributed to specific cell-types within the spleen, we developed Celestial, a machine-learning framework to identify cell-type-specific changes in bulk tissue samples. We found that age-related bulk molecular changes could be attributed in part to compositional remodeling of cell-types—expansion of GZMK+ CD8+ T cells and C1Q+ macrophages alongside naive T cell and global B cell loss. These results demonstrate that cell-type-aware interpretation can inform bulk multi-omic data for accurate mechanistic inference in heterogeneous tissues undergoing complex molecular remodeling.

## Introduction

Aging is a complex, temporal process marked by gradual decline of organ function, increasing frailty, and deteriorating health. Multi-omics approaches have advanced our understanding of aging by revealing molecular underpinnings of these changes^1–5^. Proteomic, metabolomic, and lipidomic data further demonstrate that chronic inflammation and oxidative stress disrupt protein homeostasis, leading to functional changes and damage to proteins and lipids^6–8^. Proteomic signatures consistently highlight immune and inflammatory pathways as major contributors to age-related diseases, underscoring immune dysfunction as a key driver of aging pace and quality^2,9–12^.

A fundamental challenge in aging biology is the poor correlation between mRNA and protein abundances, even at steady state, due to post-transcriptional regulation of mRNA stability, translational efficiency, and protein half-life^13–15^. Multi-omics studies of tissues across multiple organs have consistently shown that protein and RNA levels frequently change independently of age, revealing an aging-associated uncoupling of the transcriptome and proteome that is not detectable by either modality alone^16,17^. RNA-protein abundance uncoupling implies that post-transcriptional mechanisms, altered mRNA stability, translational reprogramming, and changes in protein turnover, contribute to the aging proteome independently of transcriptome^12^. Understanding when and how this uncoupling arises is essential for interpreting molecular aging data.

Single-cell transcriptomics adds another layer of insight into the aging process by revealing cellular heterogeneity and age-related changes in cell populations such as stem-cell depletion, inflammatory cell infiltration, and accumulation of senescent cell populations^18–21^. The Tabula Muris Senis (TMS) atlas illustrates that bulk tissue changes often reflect altered cellular composition rather than uniform aging within cell types^19^. While single-cell transcriptomics can profile up to 100,000 cells per experiment, single-cell proteomics remains limited in scale^22,23^. Antibody-based mass cytometry can quantify up to 30–50 proteins across millions of cells, offering low proteome coverage across many cells^24,25^. Mass spectrometry based single-cell proteomics improves coverage up to ∼800–5,000 proteins^26–28^, but current datasets are limited to a few thousands individual cells^22,23^. Thus, at present, indirect computational approaches are needed to integrate transcriptome and proteome data at the single-cell level.

The spleen, a central immune organ integrating innate and adaptive functions, is well suited for studying immune aging. It contains diverse immune cell types, with lymphocytes as the dominant population. Naive and stem-like T cells expressing *Tcf7*, *Lef1*, and *Satb1* decrease, replaced by effector and exhausted T cells over time^29,30^. Additionally, macrophages are upregulated with age, the main source of complement system components coinciding with increased complement activation contributing to inflammaging^31,32^. In addition, differences between sexes are among the major biological variables in immune aging, responsible for both cell-specific and molecular alterations^33,34^.

Proteomic analyses of aging spleen have revealed progressive dysregulation of immune pathways, protein homeostasis (proteostasis), and metabolism^16,17^. Declining ubiquitin-proteasome system activity with age is a conserved feature across organisms, contributing to accumulation of damaged proteins and loss of proteostasis^35–37^. Additional proteostasis-related effects are seen in proteins involved in ER function, protein trafficking, and metabolism^16,38^. Yet, proteomics studies often end at middle age and miss later life intervals when the most significant proteomic changes occur^38^. Time-point selection is therefore critical: young adult mice (1-3 months) are still maturing, while aging trajectories are nonlinear, with major dysregulation appearing after 15-18 months. Though these samples can be challenging to acquire, comprehensive studies should include mature adults (>6 months) through geriatric ages (24–30 months) with intermediate time points to capture these nonlinear dynamics.

An unresolved challenge in tissue aging research is distinguishing whether bulk molecular changes reflect altered gene regulation within cells, shifts in the abundance of specific cell populations, or both. To address this, we profiled matched proteomes and transcriptomes from murine spleen across eight time points spanning 6 to 30 months of age in both sexes—a temporal resolution sufficient to capture progressive linear changes with age, as well as acute abundance changes. To move beyond bulk-level description and resolve the cellular origins of these changes, we developed Celestial, a computational method that integrates single-cell transcriptomic reference data with bulk multi-omics measurements to assign molecular changes to specific cell-types or globally expressed programs. Existing computational deconvolution approaches, including CIBERSORTx^39^, MuSiC^40^, and BisqueRNA^41^, estimate cell-type proportions from bulk expression data but do not resolve which genes drive observed bulk abundance changes or whether those changes arise from altered cell numbers, per-cell expression shifts, or both. Celestial addresses a complementary problem: rather than estimating how much of each cell type is present, it assigns individual genes to their most likely cell-type origin and, by integrating single-cell aging dynamics, infers the mechanistic source of each gene’s age-related bulk abundance change. Together, these approaches reveal how transcriptional, post-transcriptional, and compositional mechanisms jointly drive splenic immune aging in a sex-dependent manner

## Results

### High temporal resolution of spleen aging reveals linear and nonlinear aging patterns

We profiled spleen tissue from healthy aging C57BL/6 mice of both sexes across eight time points spanning adulthood to advanced age (6 to 30 months of age), with ∼5 animals per sex per time point (n = 76 samples; **Fig. 1A, S1A**). Matched proteomics and transcriptomics generated from the same samples yielded 9,095 protein groups and 16,719 transcripts; after matching genes across modalities, 8,528 were retained for multi-omic analysis (**Table S1-2**, **S2A**).

**Figure 1.**
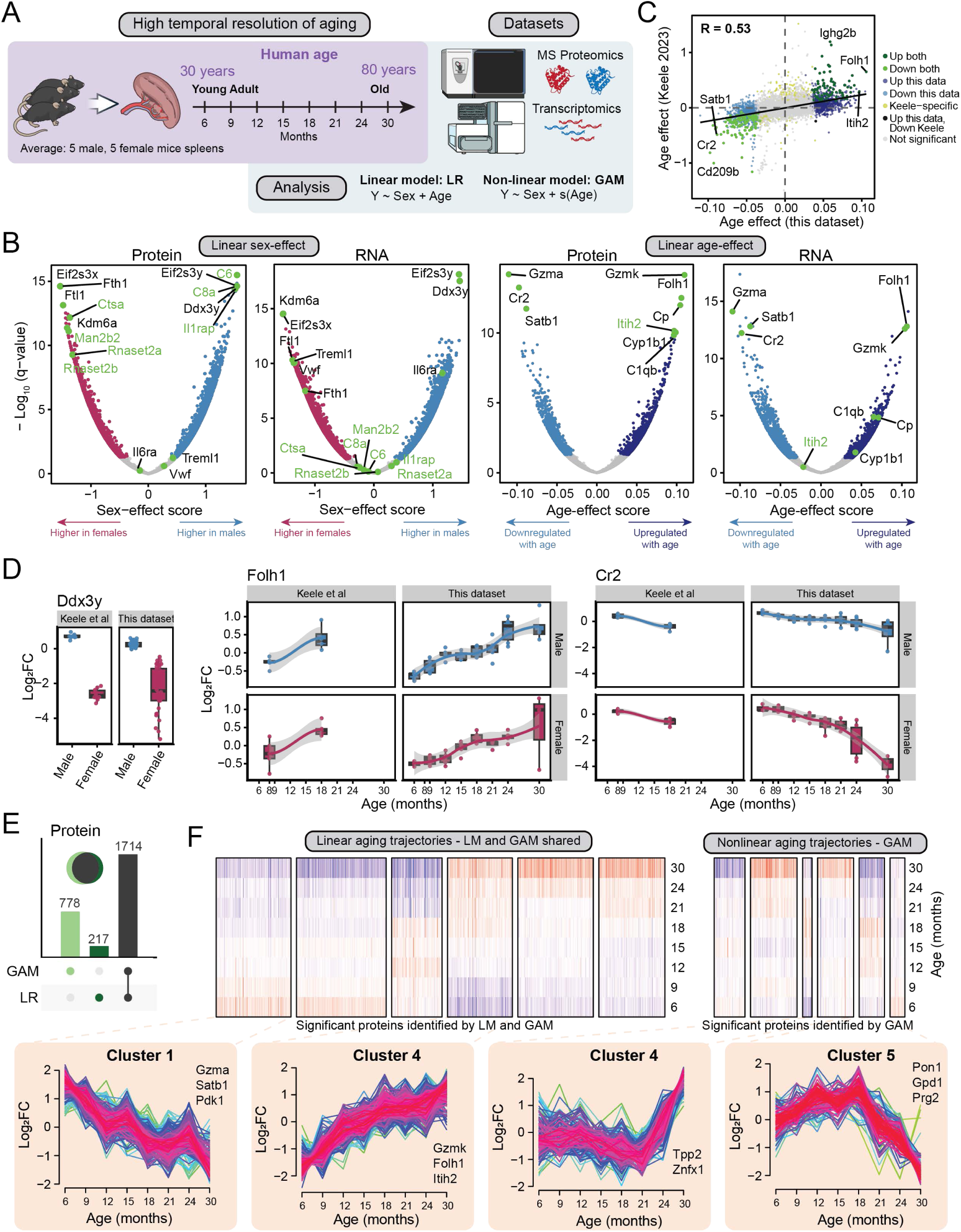
Multi-omics profiling of murine spleen aging. **A)** Experimental design: matched proteomics and transcriptomics were generated from spleens from C57BL/6 mice of both sexes (n = 76; ∼5 animals per sex per age) at eight time points, from 6 to 30 months. A total of 9,095 proteins and 16,719 transcripts were quantified; 8,528 genes were matched across both modalities and used for all subsequent analyses. Linear regression (LR) and generalized additive models (GAM) were applied to model age- and sex-related abundance changes. q-value < 0.05 indicates significant changes. **B)** Volcano plots of protein and RNA sex- and age-effects, showing linear regression coefficients across all 8,528 genes. Selected genes with the largest sex- and age-associated changes are labeled. **C)** Scatter plot comparing protein age-effect scores between this study and Keele et al. 2023 spleen data (8 and 18 months). r_Spearman_=0.53. Selected genes of interest are labeled. **D)** Plots comparing abundance change of DDX3Y, FOLH1 and CR2 proteins in this study and Keele et al. 2023, highlighting comparable age-associated trends. Data from this dataset: log2 fold change of abundance of gene / median abundance across all samples, Keele et al dataset: log2 abundance of gene / pooled sample. **E)** Upset plots showing overlap of proteins significantly changed with age by LR (1,931) and GAM (2,492), with 1,714 identified by both methods (q-value < 0.05). **F)** Heatmaps of shared age-regulated proteins identified by LR and GAM indicating linear trends in abundance change, as well as GAM-identified nonlinear proteins, clustered by temporal expression pattern. Clusters highlight proteins with pronounced linear changes on left, and nonlinear change in abundance on right, with peaks at 18 and 21 months of age.

Principal component analysis (PCA) of proteomics data showed no batch-related variability (**Fig. S1B**), and we observed no batch-dependent differences in protein–RNA sample correlations (**Fig. S1C**). As expected, sex was the primary driver of variance (PC1) in both proteomics and transcriptomics datasets (**Fig. S1B**). Age explained less overall variance than biological sex (PC3; **Fig. S1B**), consistent with previous findings^16,17^. Sex-associated differences were confirmed at both the protein and RNA levels, with male-biased expression of Y chromosome genes *Ddx3y* and *Eif2s3y* and female-biased expression of *Eif2s3x* and *Kdm6a* (**Fig. 1B, S1G**). Sex-dependent molecular variability became increasingly pronounced with age, as described below. While we explore this more in the next section, the per-sample correlation between protein and RNA abundances remained stable during early adulthood and increased significantly at ages 24 and 30, driven largely by consistently higher correlations in female samples (**Fig. S1D,E**).

The high temporal resolution of our aged murine samples enabled us to detect both linear and nonlinear changes in protein abundance with age. To do this, we applied linear regression (LR) and generalized additive model (GAM, non-linear) analyses to model age-related trajectories (**Table S1**). We identified 1,931 and 2,492 proteins with significant age-associated abundance changes by LR and GAM, respectively, with 1,714 detected by both methods (**Fig. 1E**). Among proteins with significant linear age effects, granzymes A (GZMA) and K (GZMK) showed some of the strongest changes, with opposing age-associated abundance patterns (**Fig. 1B**). Additional proteins upregulated with age included FOLH1, CP, ITIH2, and CYP1B1, while CR2 and SATB1 were among the most significantly downregulated. These findings were consistent with aging spleen data from Keele et al. spanning 8 and 18 months^16^, with concordant age- and sex-related protein effects (Spearman r = 0.53 and r = 0.47; **Fig. 1C-D, S1F**).

Because GAMs capture nonlinear age trajectories that linear models may fail to detect, the two modeling approaches were sensitive to different patterns of age-associated change (**Fig. 1E, S1I-K**). Clustering of shared and GAM-unique age-regulated proteins revealed distinct temporal patterns, with subtle abundance shifts at 12–15 months and more pronounced changes at 18 and 21–24 months (**Fig. 1F**). These time points correspond to midlife and early elderly stages in humans (∼45 and ∼60 years)^2,42^, underscoring the importance of sampling across the full adult lifespan to capture the nonlinear dynamics of molecular aging that would be missed by studies ending at middle age.

### Protein and RNA abundances reveal concordant and discordant age-related changes

Transcriptomic and proteomic measurements from the same samples provide an opportunity to directly compare how RNA and protein levels change with age. Coupling between RNA and proteins predominantly increased throughout the lifespan. As shown previously, coregulated (significantly correlated) protein and RNA abundances can be used to identify functionally related processes^43,44^. Across samples, protein–protein correlations remained stable with age, whereas RNA–RNA correlations declined from ∼18 months onward, suggesting that transcriptional noise accumulates earlier than proteomic dysregulation (**Fig. 2A**). Consistent with this, proteins slightly outperformed RNAs in predicting chronological age using elastic-net linear regression (protein: R² = 0.876, MAE = 8.39; RNA: R² = 0.838, MAE = 8.99; **Fig. 2B, S2B**), indicating that the two modalities carry partially non-overlapping information about biological age. Age-related reduction of RNA-RNA but not protein-protein correlation suggests that transcriptional noise accumulation^45–47^ may precede changes in the proteome and that post-transcriptional buffering initially absorbs transcriptional dysregulation before starting to fail^12,15^.

**Figure 2.**
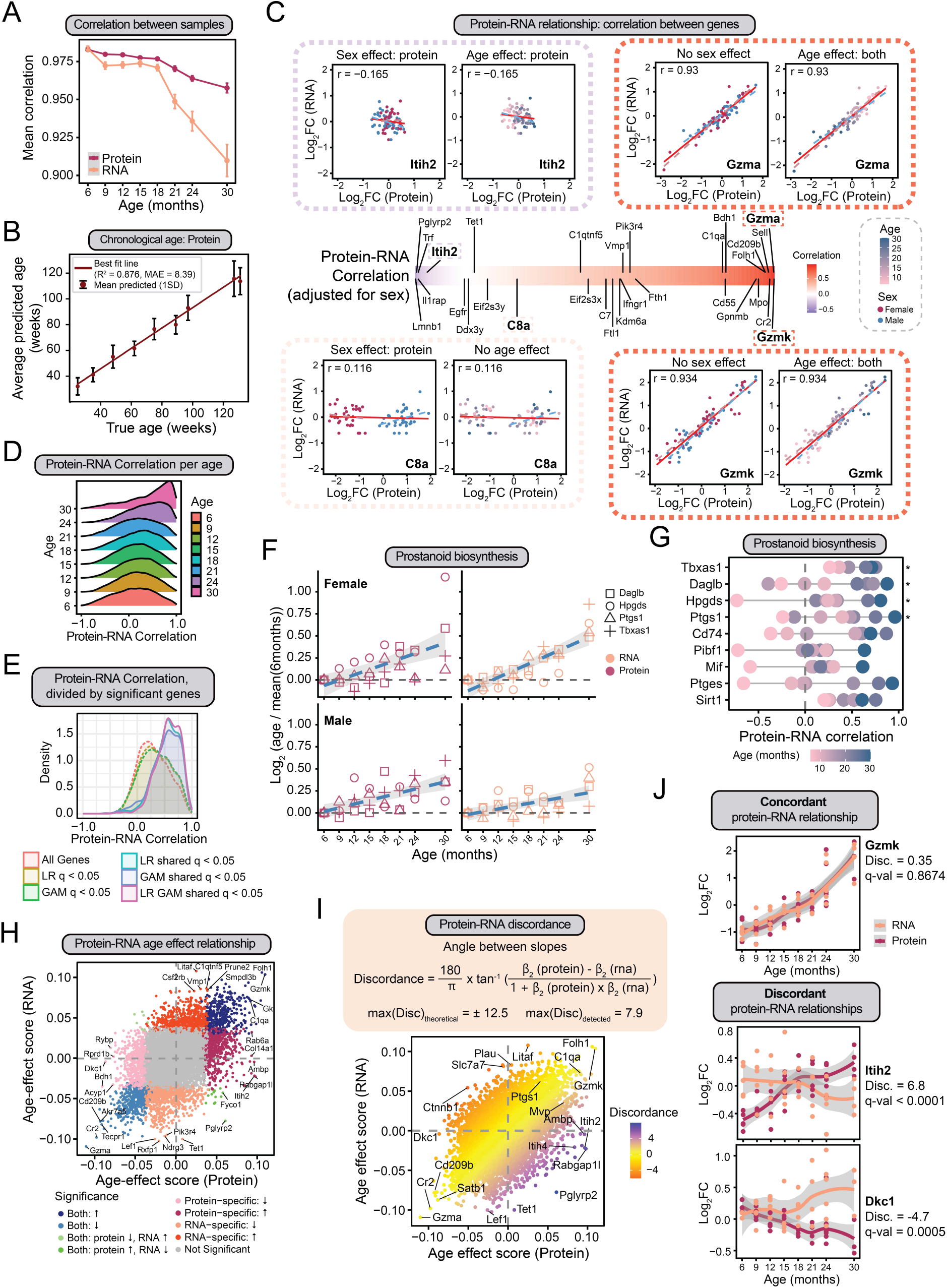
Protein–RNA relationship dynamics across the murine spleen lifespan. **A)** Sample-level pairwise Pearson correlations within proteomics (protein–protein) and within transcriptomics (RNA–RNA) datasets across all ages. Lines indicate mean per age with standard deviation. **B)** Chronological age prediction from protein data using elastic-net linear regression. Scatter plots show predicted versus actual age. Model performance: protein R² = 0.876, MAE = 8.39 months. **C)** Per-gene partial correlations between protein and RNA abundance across all samples (adjusted for sex). Individual plots show genes with high protein-RNA correlation (Gzma and Gzmk), and low protein-RNA correlation (Itih2 and C8a), colored by age and sex. **D)** Distribution of per-gene partial correlations between protein and RNA abundance adjusted for sex, calculated separately at each age. **E)** Density plots showing distributions of protein–RNA partial correlations for gene subsets: genes significant in both modalities by LR, GAM and shared between LR and GAM (full lines; r_median(LRshared)_ = 0.56, r_median(GAMshared)_ = 0.56, r_median(LR-GAMshared)_ = 0.58), and all significant genes in either modality by LR or GAM and all genes in our data (dashed lines; r_median(All)_ = 0.28, r_median(LR q<0.05)_ = 0.32, r_median(GAM q<0.05)_ = 0.34). **F)** Protein abundance and RNA trajectories across age for four prostaglandin biosynthesis genes with significant age effects on protein and RNA level: Tbxas1, Hpgds, Ptgs1 and Daglb. **G)** Protein–RNA partial correlations for proteins in prostaglandin pathway, showing correlation values per age. Significant genes are indicated with an asterisk. **H)** Scatter plot of protein versus RNA age-effect scores, with highlighted genes having significant changes on protein and RNA levels, as well as protein-specific and RNA-specific changes. **I)** Discordance between protein and RNA age-effect scores, measured as the angle between the aging trajectories of these modalities. Significance of discordant genes is calculated by Wald test, q-value < 0.05. Top protein-dominant (n = 224) and RNA-dominant (n = 195) genes are highlighted, colored in blue or red. Concordant genes are colored in yellow. **J)** Protein and RNA abundance trajectories across age for selected discordant and concordant genes: Gzmk (male), Itih2 (male), Dkc1 (female).

To examine how protein and RNA levels change together across the lifespan, we calculated per-gene partial correlations between protein and RNA abundance across all samples, adjusting for sex (**Fig. 2C**, **Table S1**), and further stratified by age (**Fig. 2D, S2C**). Genes with the gradual age-related increase in RNA-protein coupling were associated with prostanoid biosynthesis, cytoplasmic translation, RNA localization and ribonucleoprotein complex biogenesis (**Fig. S2D-E**). The overall median protein–RNA correlation across all 8,528 genes was modest (r = 0.28; **Fig. 2E**), consistent with the well-established contribution of post-transcriptional regulation to protein abundance^13–15^. Genes with significant age effects at both protein and RNA levels had substantially higher correlations (median r = 0.56–0.58) than the dataset as a whole, suggesting that transcriptional regulation contributes more directly to protein abundance for these genes^13^. However, many genes changed significantly at only one level, diluting overall coregulation and pointing to widespread regulation beyond transcription.

### Concordant protein-RNA associations reveal dysregulation of anti-inflammatory pathways in aging spleens

Among the genes with increased protein-RNA correlation in prostanoid synthesis, four out of nine had significant increases in protein abundances with age (**Fig. 2F).** Increase in the correlation of protein and RNA abundances of the prostaglandin synthetases *Tbxas1*, *Hpgds*, and *Ptgs1* (*Tbxas1*: r_6month_ = 0.26; r_30month_ = 0.71; *Hpgds:* r_6month_ = −0.73; r_30month_ = 0.81; *Ptgs1:* r_6month_ = −0.24; r_30month_ = 0.96; **Fig. 2F-G**) and increase in their protein abundances throughout age suggest that transcriptional regulation of these specific synthetases later in life may be the contributing mechanism to increased prostaglandin production with age, which was previously shown to drive reduction in T cell function^48^. The protein–RNA correlation of the major vault protein, Mvp, a member of the large ribonucleoprotein vault complexes that play a role in RNA localization, increased with age (r_6month_ = −0.05; r_30month_ = 0.80, **Fig. S2F**). The significant increase in MVP protein abundance, coupled with a non-significant change in transcript levels and an age-dependent increase in protein–RNA correlation, suggests that Mvp accumulation in older mice may be driven by a transcriptional program—potentially as a compensatory mechanism to reinforce Mvp’s anti-obesity and anti-inflammatory roles during splenic aging^49,50^.

### Discordant protein-RNA associations reveal dysregulation of telomerase and protostasis

To investigate potential regulatory mechanisms beyond transcriptional programs, we quantified RNA-protein discordance, a divergence in the direction or magnitude of age-related abundance changes between the two modalities—a distinct concept from RNA-protein correlation, which reflects steady-state coupling of expression levels. We measured discordance as the angle between protein and RNA age-effect slopes for each gene, where a discordance angle of zero indicates perfectly concordant change and larger angles indicate increasingly divergent age-related trajectories (**Fig. 2H-I, Table S1**; see Methods). Among concordant genes—those changing in the same direction and magnitude at both levels—were immune effectors *Gzmk*, *Gzma*, *C1qa*, *Folh1*, and *Cr2* (**Fig. 2I**). *Gzmk* showed concordant increases for both protein and RNA abundances with age, consistent with clonal expansion of *Gzmk*⁺ CD8⁺ T cells and suggesting a cell-intrinsic transcriptional response to aging^24^ (**Fig. 2I-J**). In contrast, a total of 419 genes had significant RNA-protein discordance (q-value < 0.05). *Itih2* and *Itih4*, protease inhibitors that crosslink hyaluronan in the extracellular matrix, increased substantially at the protein level with no corresponding RNA change, suggesting post-translational stabilization or reduced protein clearance (**Fig. 2H-J**). Conversely, *Dkc1*, a pseudouridine synthase component of the telomerase holoenzyme, decreased at the protein level without a corresponding transcript change, pointing to impaired complex assembly or accelerated protein degradation with age^51^ (**Fig. 2J**). Among the genes with significant RNA-protein discordance is *Lef1*—a naive T cell transcription factor (**Fig. 2I**). *Lef1* RNA was among the most significantly downregulated with age in bulk data, yet LEF1 protein abundance remained stable across the entire lifespan (**Fig. 2I**). As we show later, this bulk-level observation reflects loss of the *Lef1*⁺ T cell population rather than transcriptional downregulation within expressing cells.

### Pathway and protein complex dynamics reveal coordinated age-related remodeling

Many of the pathways with RNA-protein discordance identified above are mediated by cellular pathways or protein complexes rather than individual proteins^16,52^. Thus, pathway-level and complex-level analyses provide an additional layer of resolution for discerning age-related changes in the proteome. Gene set enrichment analysis (GSEA) of protein age-effect scores revealed a broad upregulation of inflammatory and hemostatic programs—including platelet activation, collagen fibril organization, innate immunity, and complement activation—consistent with a pro-inflammatory proteomic shift during splenic aging (**Fig. 3A**, **Table S2**). Endoplasmic reticulum and Golgi-associated pathways were similarly elevated, whereas RNA-related processes such as mRNA processing, splicing, and cytoplasmic translation were downregulated at the protein level. To further dissect the directionality of RNA–protein discordance, we compared relative age-associated changes across protein and RNA layers. Genes whose protein abundance increased disproportionately relative to their transcript levels were enriched for proteostasis-related functions, including cytoplasmic translation and ER maintenance, as well as adaptive immunity. Conversely, genes with disproportionately greater transcript increases were enriched for telomere maintenance and mRNA processing (**Fig. 3A**, **S3A-B**). Notably, mRNA processing and splicing machinery showed no significant downregulation at the RNA level, indicating that their loss observed at the protein level is driven by post-transcriptional mechanisms rather than reduced transcription. The divergence between protein and RNA pathway enrichments confirms that the two modalities capture distinct aspects of splenic aging that are not interchangeable.

**Figure 3.**
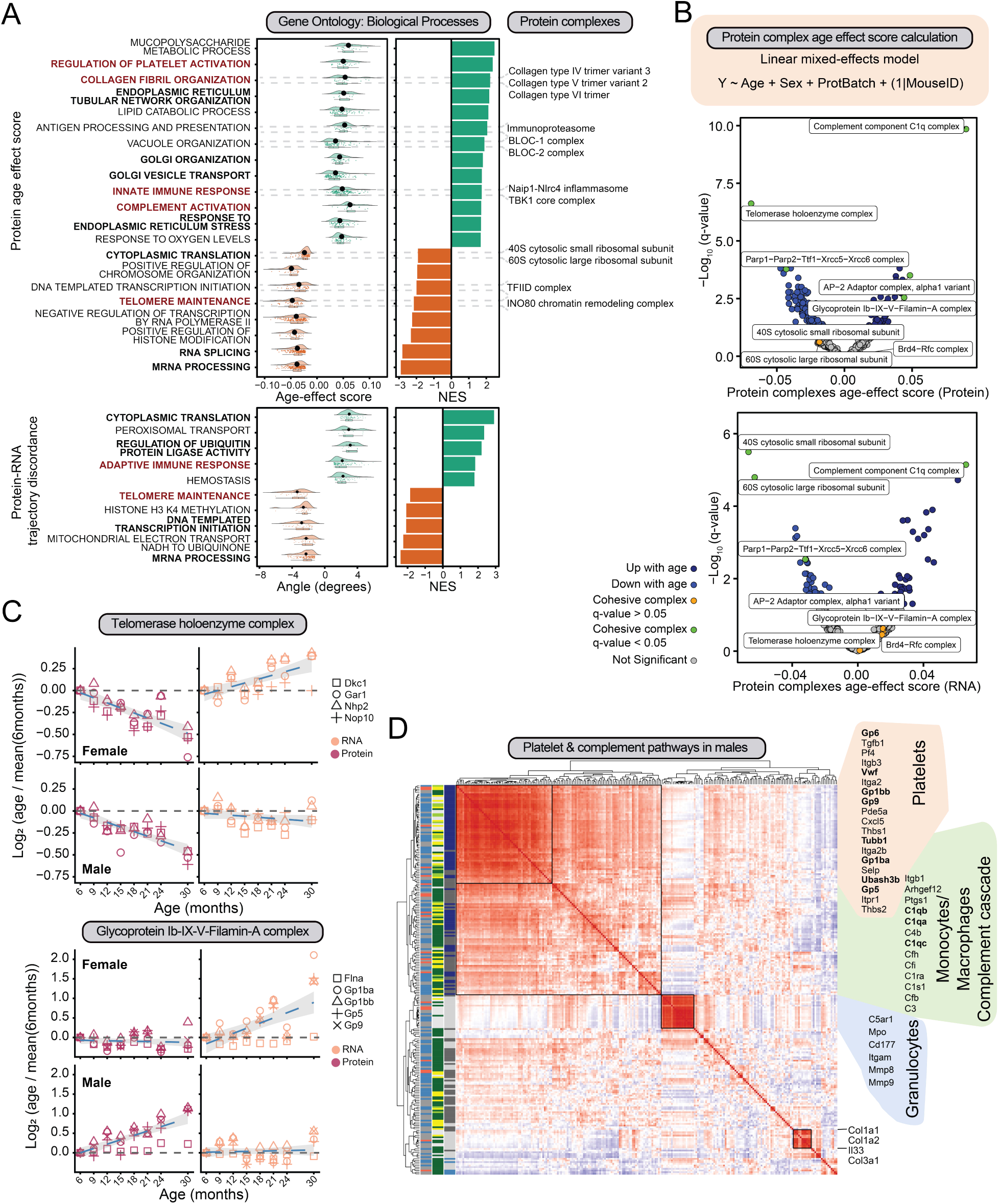
Pathway-level and protein complex dynamics during splenic aging. **A)** GSEA results for protein age-effect scores (coefficient) and protein–RNA discordance (angle). Normalized enrichment scores (NES) are shown for selected GO biological pathways (right) and distribution of age-effect score values (top) or angle values (bottom) for genes in the leading edges. Positive NES indicates enrichment among increasing or protein-dominant genes; negative NES indicates enrichment among decreasing or RNA-dominant genes. Protein complexes that belong to the biological pathways are indicated on the right. **B)** Analysis of protein complexes using both modalities using linear mixed-effects models. Shown are volcano plots of protein complexes with significant age effects on the protein or RNA level (q-value < 0.05) and high median pairwise partial correlation among subunits (cohesiveness, green), or highly cohesive protein complexes not changing with age (orange). **C)** Protein and RNA abundance trajectories across age for four members of telomerase holoenzyme complex (top) and five members of Glycoprotein Ib–IX–V–Filamin-A complex (bottom). **D)** Correlation matrix of proteins belonging to platelet, complement and coagulation pathways in males, highlighting distribution of correlation between platelet markers, macrophage markers and complement proteins, and granulocyte proteins.

Since protein complex members tend to be coregulated at the protein level to preserve stoichiometry and function ^16,52^, we examined complex-level dynamics across age by combining CORUM and Complex Portal annotations^53,54^ and estimating complex-level age effects using mixed linear models; cohesiveness was measured as median inter-member partial correlation (**Fig. 3B**, **Table S3**). The most significantly changing highly cohesive complexes were the Complement C1q and Telomerase holoenzyme (**Fig. 3B–C**). Aging was associated with coordinated loss of Telomerase holoenzyme proteins without a corresponding change in RNA, implying post-transcriptional loss of function consistent with impaired complex assembly or reduced stoichiometry^51,55^. RNA downregulation of 40S and 60S cytosolic ribosomal subunits without corresponding protein changes suggests translational buffering, consistent with the long half-life of ribosomal proteins within assembled complexes^56^ (**Fig. 3B**, **S3C–D**). Additional complexes with protein-specific age effects included the Glycoprotein Ib–IX–V–Filamin-A and AP-2 Adaptor complexes. Together, these findings demonstrate that pathway and complex-level analyses reveal coordinated age-related changes—and RNA-protein discordances—that are not apparent from individual gene analysis alone.

### A male-specific platelet-macrophage inflammatory axis emerges with age

Among the most significantly upregulated cohesive complexes were complement complexes C1q and C4b2a3b-B (classical/lectin C5 convertase), as well as the platelet-associated complexes Glycoprotein Ib–IX–V–Filamin-A and *Itga9*–*Itgb1*–*Thbs1*. Increased abundance of members of these complexes was consistent with pathway-level enrichment for platelet activation and complement activation described above (**Fig. 3B–D, S3B–C, Table S3**). In general, coregulated complex proteins also had concordant protein–RNA abundances with age (**Fig. S3C**). Consistent with aging-associated expansion of macrophage C1q production, the shared transcriptional coregulation of all three subunits of the complex (*C1qa*, *C1qb*, and *C1qc*) in macrophages ^32,57^ may lead to coupling between RNA and protein levels during aging. Indeed, we observed an age-dependent increase in protein–RNA correlation for C1qb (r_6month_ = 0.00; r_30month_ = 0.79) (**Fig. S2F**). We noted that this upregulation was especially pronounced in males. Upregulation of *Tbxas1*, through production of thromboxane A2, suggests increased platelet activation and aggregation with age^58,59^. In addition, activated platelets secrete CXCL5, which acts as a chemoattractant for monocytes and neutrophils^60^; consistent with this, CXCL5 protein abundance increased with age in males but not females, with no change in transcript abundances in either sex (**Fig. S1H**). Male-biased platelet proteins correlated strongly with macrophage and complement proteins, but not with granulocyte proteins (**Fig. 3D**), suggesting platelet–macrophage crosstalk^61^ rather than platelet–granulocyte coupling^62^ in the aging spleen. Together, these findings point to the emergence of a coordinated platelet-macrophage inflammatory axis in aging males. The specificity of this axis would be consistent with the known age-related emergence of hyperreactive platelet populations in males driven by a distinct megakaryocyte differentiation pathway^59^. Taken together, these data point to the importance of understanding the spleen is a cellularly heterogeneous organ, for which bulk measurements alone cannot determine if these discordant patterns reflect cell-intrinsic changes in gene regulation, shifts in the abundance of specific cell populations, or both.

### Cell type-specific and global aging signatures identified by Celestial

To address the question posed in the previous section—whether RNA-protein discordance in bulk molecular changes reflects cell-intrinsic regulation or shifts in cellular composition—we developed Celestial, a computational framework that integrates single-cell transcriptomic reference data with bulk multi-omics measurements to assign transcripts and proteins to specific cell types or globally expressed programs (see Methods). Several of the most significantly changing proteins in our dataset are known to be highly cell-type specific, including GZMA (NK cells), GZMK (T cells), CR2 (B cells), and C1QA (macrophages) (**Fig. 1B**) ^24,31,63^. We confirmed this based on a recent single-cell transcriptomics atlas, Tabula Muris Senis (TMS), e.g., *Cr2* in B cells and *Gzma* in NK cells, as well as global expression of the ribosomal *Rps15a* (**Fig. 4A**) ^19^.

**Figure 4.**
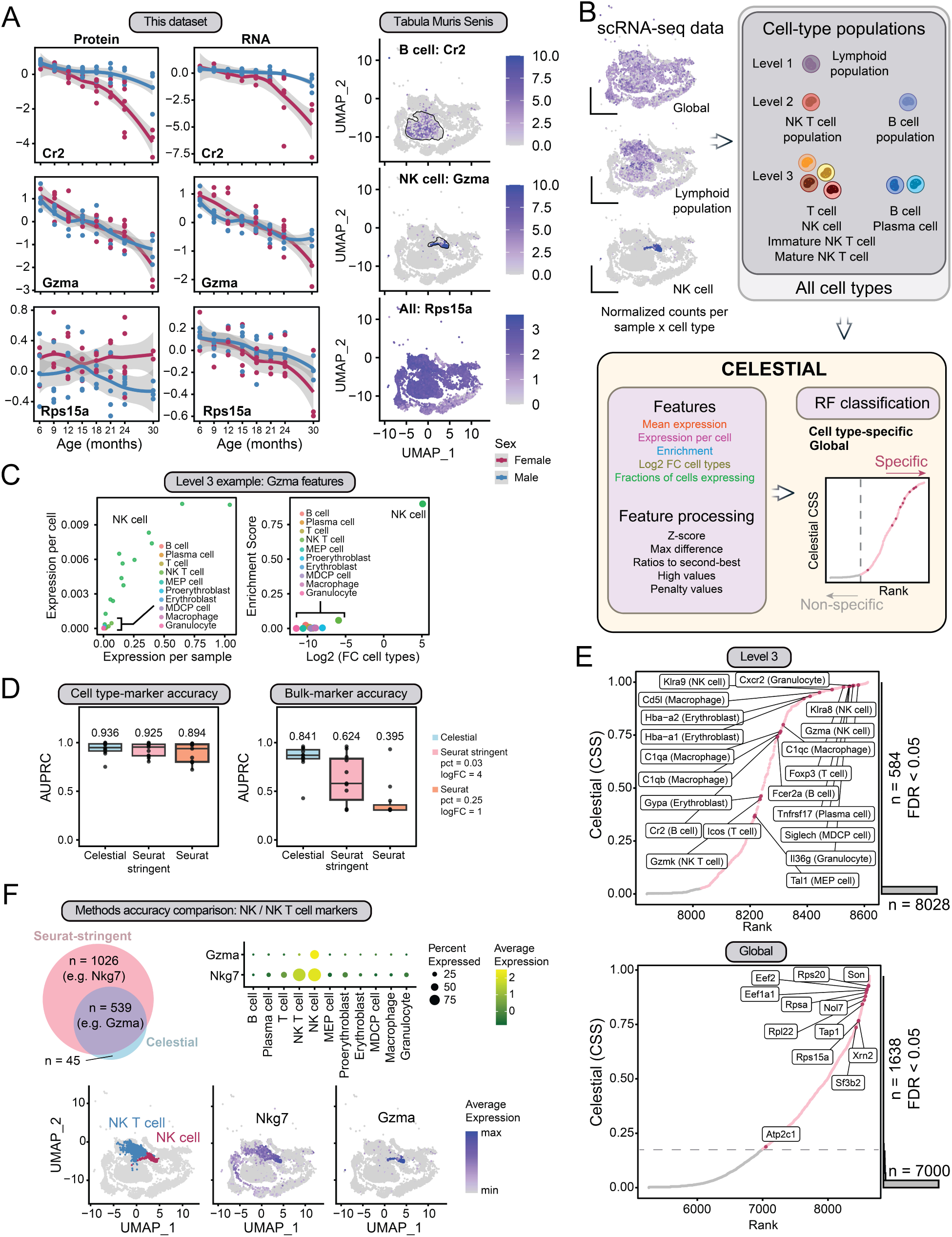
Development and benchmarking of the Celestial cell type specificity tool. **A)** Change Single-cell RNA-seq expression from TMS spleen data illustrating cell type-specific expression of Cr2 (B cell) and Gzma (NK cell), and globally expressed Rps15a, together with protein and RNA abundance trajectories across age. Cell-types are annotated using TMS metadata. **B)** Schematic of the Celestial framework: single-cell RNA-seq features are computed per gene across cell types and used as input to a random forest classifier that assigns a cell type-specificity score. Cell types are grouped in levels based on their biological characteristics into Level-3 (cell type-specific), Level-2 (the closest cell populations), Level-1 (cell families), and globally expressed genes into Level-0. Statistical significance was calculated using b-distribution. **C)** Plots showing features distributions used by Celestial for cell-type-specific gene classification. Shown are plots for Gzma, indicating specificity of Gzma expression in NK cells. Expression per cell and per sample are calculated per donor_id from TMS data, enrichment score and log2 fold change difference between cell types is calculated after averaging values per donor_id. **D)** AUPRC benchmarking of Celestial against two Seurat marker calling settings (default: pct = 0.25, logfc.threshold = 1; stringent: pct = 0.03, logfc.threshold = 4;), evaluating the presence of a gene in a cell type (TMS annotations) at single-cell, cell-type marker level (log-normalized expression) and bulk cell-type marker level (pseudobulk CPM). **E)** Overlap of Celestial-identified cell-type-specific genes at Levels 2 and 3, with known markers from the CellMarker and Descartes databases, shown per cell type, detected using EnrichR (adj p-value < 0.05). **F)** Examples of Celestial and/or Seurat identified markers, showing accuracy for Gzma as a marker in NK cells detected by Celestial and Seurat (shown for stringent parameters). Seurat inefficiently assigned Nkg7 to two cell types, NK and NK T cells.

Celestial extends the inference of canonical deconvolution methods^64^ for estimation of cell-type proportions to instead focus on gene-level cell-type assignment. Celestial uses cell-type annotations and scRNA-seq expression data from atlases such as TMS (**Fig. 4B**) to identify cell-type markers that are enriched within a cell-type and specific to a cell-type based on stringently filtered scRNA-seq transcript features. These features include mean expression, expression-per-cell, and relative enrichment of expression compared to all other cell types (**Fig. 4C**). Importantly, selection of these features enabled Celestial to extensibly identify cell-type markers across multiple tissues, including those with diverse cell-types such as spleen and kidney (**Fig. S4B-C)**. From TMS splenic annotations, we grouped cell-type populations into three resolution levels: Level-3 (11 distinct cell types: B cells, plasma cells, T cells, natural killer (NK) T cells, NK cells, megakaryocyte-erythroid progenitor (MEP) cells, proerythroblasts, erythroblasts, macrophage dendritic cell progenitor (MDCP) cells, macrophages, and granulocytes), Level-2 (4 populations: B cell populations, NK T cell populations, megakaryocyte/erythroid populations (MEP), and granulocyte/myeloid populations (GMP)), and Level-1 (2 lineages: common lymphoid populations (CLP) and common myeloid populations (CMP)). Celestial then uses a random forest classifier to calculate a cell-specificity score (CSS) and significance of whether a transcript (or its resulting protein) was specific to a cell type or globally expressed.

For canonical single-cell-level marker calling benchmarked against Seurat-derived markers^64^, Celestial had modestly improved performance compared to Seurat for single-cell-level marker calling (mean area under the precision-recall curve [AUPRC]; mean AUPRC_Celsetial_ = 0.936, AUPRC_Seurat_ = 0.894), and the difference was further reduced using a more stringent set of filtering criteria in Seurat (mean AUPRC_Seurat-stringent_ = 0.925). However, Celestial outperformed both standard and stringent Seurat settings at the bulk level (mean AUPRC_Celsetial_ = 0.841, AUPRC_Seurat-stringent_ = 0.624, AUPRC_Seurat_ = 0.395; **Fig. 4D**). The greatest performance advantage was observed for biologically related and numerically small populations such as T, NK and NK T cells or B and plasma cells (**Fig. S4D-E**). Seurat identified more overall markers, while assigning one gene to several cell types. For example, all three tested methods accurately identified Gzma as an NK cell marker (**Fig. 4F**). However, Seurat called *Nkg7* as cell-type specific for both NK cells and NK T cells, while Celestial did not call *Nkg7* as cell type specific for the analysis of bulk-omics data.

Celestial was designed to overcome marker-cell-type specificity issues, leading to better performance in assigning markers for hard to differentiate cell types, e.g., B and plasma cells, or NK T cells versus T cells and NK cells. Celestial-assigned, cell-type-specific genes (Level-3) included known cell type markers such as *Gzmk* (NK T cell in TMS annotations, corresponding to GZMK+ CD8+ T cells described by Mogilenko et al ^24^), *Foxp3* (T cell), and *Tnfrsf17* (BCMA, plasma cell) (**Fig. 4E**). Notably, Celestial was also able to assign ‘globally expressed’ genes including ribosomal proteins *Rpl22* and *Rps20*, and translation-related genes *Eef2* and *Eef1a1* (**Fig. 4E**). From the CellMarker and Descartes databases^65–67^, we confirmed that the Celestial-called cell-type genes were enriched for known cell-type markers, e.g., Celestial B cell markers enriched CellMarker B cell genes and Celestial granulocyte markers enriched CellMarker neutrophil markers (**Fig. S4F**). Collectively, these benchmarks establish Celestial assignments can be used with confidence to interpret cell-type origins of bulk proteomic changes.

### Characterisation of cell-type-specific proteins in aging murine spleens with Celestial

Celestial identified 584 Level-3 cell-type genes, 995 Level-2 progenitor cell genes, 1,720 Level-1 cell-family genes, and 1,638 globally expressed genes (FDR < 0.05; **Fig. S5A**, **Table S4**). As expected for genes expressed in numerically small populations, fewer cell type-specific genes were detectable in our bulk data than globally expressed genes (∼43% versus ∼86% for Level-3 and global, respectively; **Fig. S5A**), reflecting that cell-type-specific transcripts in minor populations fall below bulk detection thresholds—a pattern that is itself informative about relative cell type abundance. Protein–RNA correlation was substantially higher for cell-type-specific genes than for globally expressed genes across all three Celestial levels (median r_Level-3_ = 0.519, r_Level-2_ = 0.567, r_Level-1_ = 0.537, versus r_Global_ = 0.219; **Fig. 5A**, **S5C**), and age-effect concordance between protein and RNA followed the same pattern. This argues that bulk protein–RNA coupling most faithfully reflects single cell-level regulation when a gene is expressed predominantly by one cell type, and that Celestial assignments can be used with confidence to interpret cell type origins of bulk proteomic changes.

**Figure 5.**
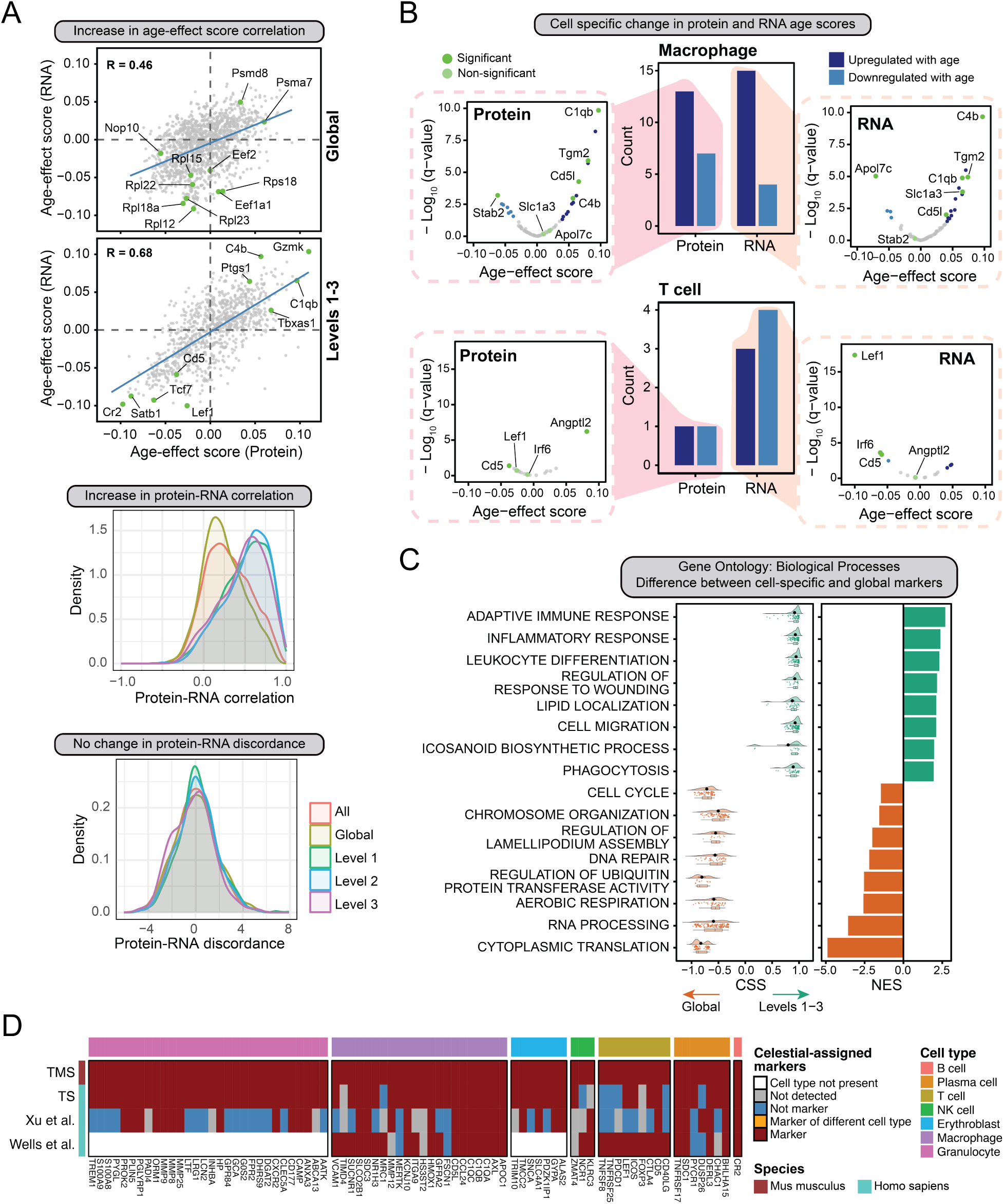
Cell type-specific aging signatures identified by Celestial. **A)** Scatter plots of protein–RNA age-effect scores correlations across Celestial specificity levels (top): cell type-specific genes across all levels r = 0.68 (Levels 1-3), and globally expressed genes r = 0.46 (Level-0), highlighting the important marker genes. Density plots of protein–RNA partial correlations (adjusted for sex, middle) and discordance (bottom) per each level. Protein-RNA correlation distributions: Level-3 r_(median)_ = 0.519, Level-2 r_(median)_ = 0.567, Level-1 r_(median)_ = 0.537, Level-0 r_(median)_ = 0.219, all Celestial genes combined r_(median)_ = 0.326. **B)** Celestial-assigned markers in macrophages and T cells: Barplots show the number of significantly changing proteins and RNAs from our aging bulk multimodal data. For each of the cell-type, volcano plots show distribution of age-effect scores versus negative log_10_ of q-values of genes assigned to be cell type specific. Highlighted genes are labeled based on their significance values (q-value < 0.05). **C)** GSEA results comparing all cell type-specific genes (Levels 1-3) versus globally expressed genes (Level-0) using Celestial specificity scores as ranking. Selected enriched pathways NES values and representative leading-edge genes CSS scores are shown for cell type-specific and global categories. **D)** Celestial-assigned markers in human scRNA-seq studies overlap with mouse TMS data. Plot shows consensus genes in cell types that exist in each of the studies^19,68–70^.

Among the cell type-specific genes quantified in our bulk data, B cell-associated genes (Level-3 and Level-2) were frequently dysregulated with age, following both linear and nonlinear patterns (53% of B cell-associated genes, **Fig. S5A-B**). Nonlinear changes were predominantly found in spleens from female mice, with proteins increasing in abundance at middle age before dropping in later life (cluster 1, 67% of nonlinear proteins; **Fig. 1F, S5D**). Proteins with linearly changing abundances largely fell into two clusters: cluster 2 proteins decreased in abundance across age (47% of proteins, female-driven) and cluster 4 proteins increased in abundance with age (32%, both sexes equally contributing). On the other hand, granulocyte and erythroblast-associated genes were upregulated with age (**Fig. S5A**). Macrophage and T cell-associated genes showed mixed patterns of up- and downregulation at both protein and RNA levels. Among macrophage-assigned markers changing linearly during aging, *C1qb*, *Tgm2*, *C4b*, and *Cd5l* changed significantly at both protein and RNA levels; *Stab2* changed at the protein level only; and *Apol7c* and *Slc1a3* changed at the RNA level only, illustrating the range of regulatory mechanisms operating within a single cell type during aging (**Fig. 5B**, **S5A**). T cell-assigned marker *Sostdc1*, Wnt and Bmp antagonist, showed nonlinear change with age. Among linearly changing T cell-assigned markers, *Cd5* decreased significantly at both protein and RNA levels, while *Lef1* had highly significant RNA-protein discordance among T cell genes with a significant decline in transcript abundance while its protein remained stable (**Fig. 5B**). Importantly, this pattern was not simply a property of the *Lef1* gene itself but reflects the behavior of a specific T cell subpopulation—a distinction we resolve using origin-of-change analysis in the following section.

To further validate the biological coherence of Celestial assignments, we performed GSEA using Celestial specificity scores, comparing cell-type-specific genes across all levels against globally expressed genes (**Fig. 5C**). Cell type-specific genes were enriched for immune processes—adaptive immunity, leukocyte differentiation, wound response regulation, and phagocytosis—whereas globally expressed genes were enriched for universally active processes including cytoplasmic translation, aerobic respiration, ubiquitin transferase regulation, and chromosome organization. Among adaptive immune response genes, we detected B cell markers (Levels 2–3) such as *Fcer2a*, *Cr2*, *Cd79a/b*, and *Cd55*, as well as T cell/NKT cell markers *Lef1* and *Pdcd1*.

Our initial Celestial cell-type assignments focused on the relevant murine TMS cell-type analyses directly useful for the murine aging proteomic and transcriptomic data presented here. In addition, we wanted to assess whether cell-type assignments could be generalized to human splenic data. Thus, we applied Celestial to human splenic scRNA-seq datasets from the Tabula Senis (TS)^68^, Xu et al.^69^ and Wells et al.^70^ (**Fig. 5D**). Cell-type marker assignments were consistent across the TMS murine and the three human scRNA-seq datasets. Cell-type marker assignments were consistent across the TMS murine and the three human scRNA-seq datasets, with every marker mapping to the same cell type in both mouse and human. The consistency of these assignments confirmed the selectivity of Celestial’s classification and the conserved transcriptional identity of major splenic cell populations between mouse and human.

### Origin-of-change of discordant RNA and protein signatures of aging

A central question raised by our bulk multi-omics analysis was whether age-related abundance changes reflected altered gene regulation within cells, shifts in the abundance of specific cell populations, or dynamics within subpopulations that alter the fraction of expressing cells. To resolve these contributions, we extended Celestial to estimate subpopulation dynamics using three metrics computed from TMS aging data for each gene and cell type: total expression, number of cells expressing the gene, and fraction of cells expressing the gene (**Fig. 6A**). Each metric was tested for age association by linear regression, allowing us to assign the cellular origin of bulk abundance changes with age.

**Figure 6.**
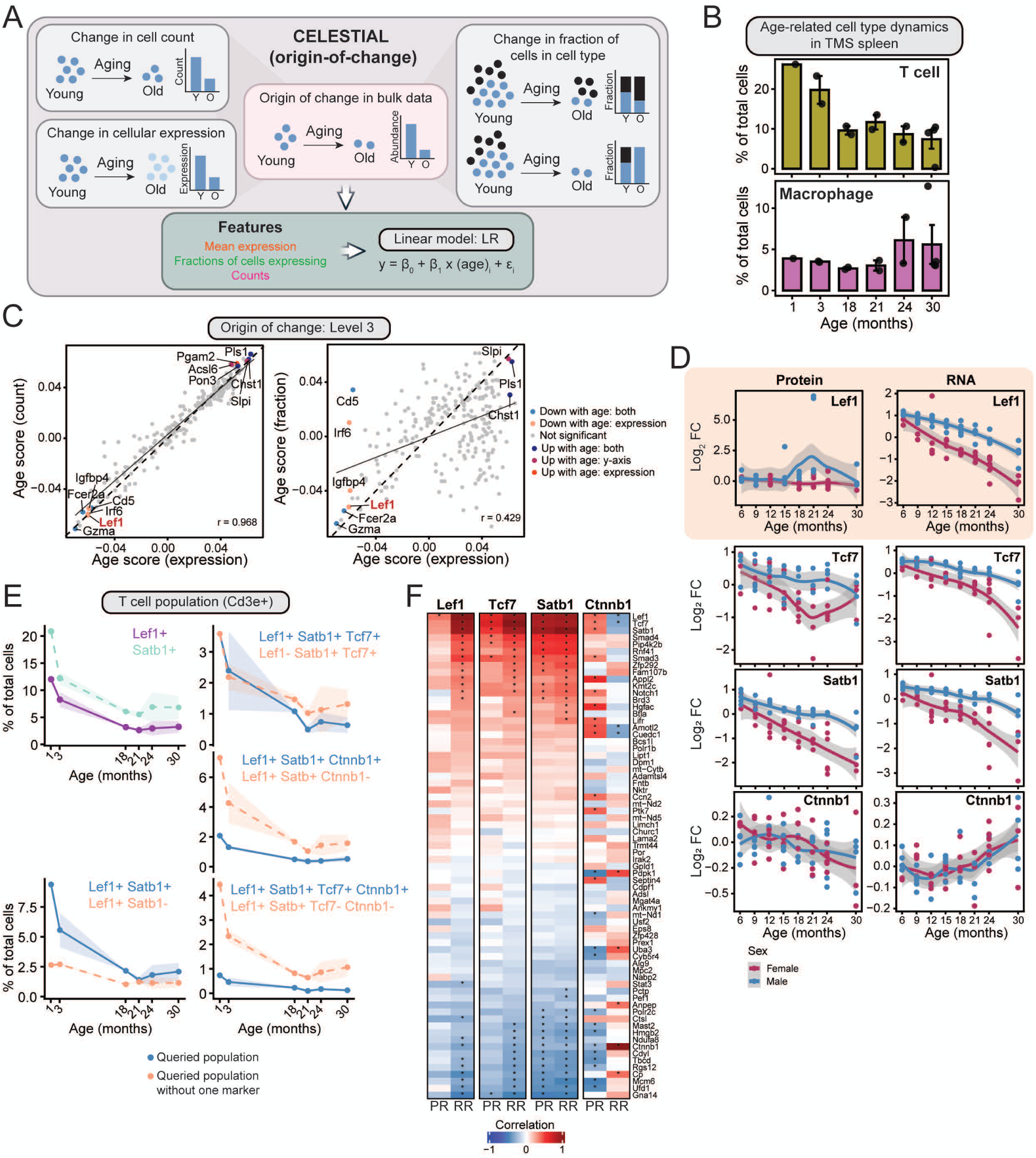
Origin-of-change analysis of age-related dynamics in specific cell types. **A)** Schematic of the Celestial origin-of-change analysis. Three feature metrics are computed from TMS aging data per gene and cell type: total expression, number of expressing cells, and fraction of cells expressing the gene. Each metric is tested for age association by linear regression, and p-values are adjusted for multiple comparisons per cell type. **B)** Cell abundance trajectories across age in TMS spleen data for T cells and macrophages, shown as percentage of total cells (cell type-specific number of cells versus total number of cells per donor_id). **C)** Origin-of-change plots, showing age-effect scores for expression, number of cells expressing the gene, and fraction of cells expressing the gene, calculated per cell type. Among the highlights genes are Gzma (all three metrics declining, consistent with NK cell loss), Cd5 (expression and cell count declining while expressing fraction increases), and Lef1 (all three metrics declining, consistent with loss of a Lef1+ T cell subpopulation). **D)** Protein and RNA abundance trajectories across age for Lef1, Tcf7, and Satb1. **E)** Trajectories of T cell subpopulations in TMS data across age, shown as percentage of total cells. Left: Lef1+ T cell and Satb1+ T cell subpopulations (Cd3e+, left); Middle: Lef1+ T cell subpopulations also positive or negative for Satb1 (Satb1+ or Satb1-), and Satb1+Tcf7+ T cell subpopulations also positive or negative for Lef1 (Lef1+ or Lef1-); Right: Lef1+Satb1+ T cell subpopulations also positive or negative for Ctnnb1 (Ctnnb1+ or Ctnnb1-) or Ctnnb1 and Tcf7 (Ctnnb1+Tcf7+ or Ctnnb1-Tcf7-). **F)** Heatmap showing partial correlation adjusted for sex of proteins and RNAs of Lef1, Tcf7, Satb1 and Ctnnb1 with RNA abundances of Wnt-target genes (MSigDB, M3: Ctnnb1 target genes). Asterisk indicates significant correlation with q-value < 0.05. PR: protein-RNA correlation; RR: RNA-RNA correlation.

In TMS, T cells declined with age whereas macrophages remained stable (**Fig. 6B**). Across cell-type-specific genes, transcript age effects on single-cell total expression and cell count were highly correlated, indicating that most age-related transcript changes in cell type-specific genes were driven by changes in cell number rather than cell-intrinsic changes in expression per cell (**Fig. S6A-B**). Age effects on the fraction of cells expressing a given transcript were more heterogeneous, suggesting that subset-specific dynamics—where some genes change within a fraction of a cell type independently of overall cell type abundance—also contribute to bulk aging signatures. As previously reported ^24,33^, *Gzma* declined with age across all three metrics—total expression, cell count, and expressing-cell fraction—consistent with loss of the GZMA⁺ NK cell population and confirming that the reduction we observed in bulk data (**Fig. 4A**) reflects a compositional shift rather than cell-intrinsic downregulation. Not all genes showed such straightforward dynamics. Among T cell-assigned genes, *Cd5* exhibited significant age-associated decreases in both total expression and expressing cell counts, yet the fraction of T cells expressing *Cd5* increased with age (**Fig. 6C**). This pattern implies complex dynamics within the T cell compartment, where the overall decline of T cells proceeds faster than the decline of the *Cd5*⁺ T cell subpopulation—a distinction that would be invisible to bulk analysis alone.

### Origin-of-change distinguishes regulatory mechanisms of *Lef1* cell-type-specific aging

The most striking and mechanistically informative example of origin-of-change resolution in our dataset was for *Lef1*. *Lef1*, along with *Tcf7*, maintain T cell identity and naive T cell survival. In our bulk transcriptomics data, *Lef1* was among the most significantly downregulated transcripts across the entire lifespan, yet LEF1 protein abundance remained stable at all ages—exhibiting significant RNA-protein discordance in our dataset (**Fig. 6D**). *Tcf7* and *Satb1* were among the highest correlating transcripts with *Lef1*, together with other known T cell and NKT cell markers, including *Tet1* and *Itk*, while no similar correlation was observed on protein level (**Fig. S6C**). Origin-of-change analysis resolved this paradox: all three metrics—total *Lef1* expression, number of *Lef1*-expressing cells, and fraction of T cells expressing *Lef1*—declined with age in TMS data (**Fig. 6C**), confirming that the RNA decline reflects loss of the *Lef1*⁺ T cell population rather than transcriptional downregulation within expressing cells. The bulk-level RNA-protein discordance observed for *Lef1* therefore arises from a compositional shift—the contraction of a specific T cell subpopulation—combined with active post-transcriptional stabilization of LEF1 protein within the surviving *Lef1*⁺ T cells.

To understand the functional context of this stabilization, we examined the identity of *Lef1*⁺ T cells and the signaling pathways in which LEF1 operates during aging. The majority of *Lef1*⁺ T cells in young mice co-expressed *Satb1*, and both populations declined in parallel with age (**Fig. 6E**). However, within the *Lef1*⁺*Satb1*⁺ population, cells co-expressing *Ctnnb1*—reflecting canonical Wnt signaling—represented only a small subset, and LEF1 protein abundance had no significant correlation with the RNA abundances of Wnt target genes (**Fig. 6F**), suggesting that LEF1 function in these cells was independent of canonical Wnt/β-catenin signaling. Instead, the maintained LEF1 protein likely supported β-catenin-independent roles in genome organization, including chromatin loop maintenance, enhanceosome assembly, and epigenetic silencing of lineage-inappropriate genes through intrinsic histone deacetylase activity. By contrast, the upstream regulators of *Lef1*— *Satb1* and *Tcf7*—declined at both protein and RNA levels with age^71–73^ (**Fig. 6D**), suggesting that the protein stabilization observed for LEF1 was specific rather than a general feature of this T cell population, and may reflect impaired proteasomal degradation or other post-translational mechanisms that selectively maintain LEF1 as this population contracts.

Together, the multi-omics profiling, Celestial cell-type assignments, and origin-of-change analysis presented here reveal that molecular aging of the spleen is shaped by the interplay of three distinct mechanisms: shifts in cellular composition that dominate the bulk signal for most cell type-specific genes; post-transcriptional regulation that maintains or suppresses protein abundance independently of transcript levels within specific cell types; and transcriptional programs that drive coordinated RNA and protein changes reflecting the intrinsic response of stable or expanding populations to the aging environment. Distinguishing these contributions—as Celestial enabled—is essential for interpreting bulk molecular aging data and for identifying the cell types and regulatory mechanisms that are most vulnerable to age-related dysfunction.

## Discussion

The spleen serves simultaneously as a site of immune surveillance, blood filtration, and hematopoietic reserve. This convergence of functions makes it a tissue where age-related changes in cellular composition, immune function, and proteostasis intersect. By profiling murine spleens across eight time points spanning 6–30 months of the murine lifespan, this study provides a temporally resolved view of how these processes unfold in parallel in matched proteomic and transcriptomic analyses. The temporal resolution of our murine aging time course allowed us to detect both gradual, linear molecular changes and more abrupt non-linear transitions—particularly at 18 months and again after 21–24 months—that would be difficult to detect in studies comparing fewer timepoints across aging^16,17^. These non-linear inflection points likely reflect discrete biological thresholds, such as the progressive loss of naive T cell homeostasis, hormone associated systemic changes in late life, or the expansion of age-associated immune cell populations, rather than simple accumulation of damage over time ^19,74,75^.

Protein-dominant changes in the ubiquitin-proteasome system, including accumulation of multiple proteasome subunits and ERAD components, point to impaired proteostasis as a mediator of splenic aging, consistent with previous work in mouse^16,17^ and human proteomes^10^. Proteostasis failure is one of the recognized hallmarks of aging^1^, and its manifestation here in an immune organ raises the question of whether impaired protein clearance contributes to the functional decline of splenic immune responses—for instance, by allowing accumulation of misfolded or damaged proteins that trigger inflammatory signaling^1^. This connection between proteostasis and immune aging remains an area of active investigation and these data identify specific molecular targets—particularly proteasome subunits and ERAD components—that may link proteostasis failure to immune dysfunction.

A key finding from the integration of RNA and protein datasets is that cell-type-specific genes consistently had stronger protein–RNA correlation and larger age-effect scores than globally expressed genes. As with previous aging studies in human tissues, overall RNA-protein correlation was relatively modest^10^. This is consistent with a model that bulk protein–RNA coupling is more tightly regulated within individual cell types than across the tissue as a whole, and that this coupling becomes more detectable when a gene’s expression is dominated by a single population. This observation has practical implications: it suggests that bulk multi-omic studies of heterogeneous tissues may systematically underestimate the strength of molecular coordination within cell types, and that deconvolution^64,76,77^ or cell-type assignment should be considered a routine step in the interpretation of such data.

A central challenge in interpreting bulk molecular data from a heterogeneous tissue is distinguishing changes in tissue composition from changes in the molecular state of individual cell types^64^. Existing deconvolution approaches estimate cell-type proportions but do not inherently seek to resolve the mechanistic origin-of-change in bulk abundance—whether it arises from altered cell numbers, altered per-cell expression, or both. Single-cell marker identification tools such as Seurat^78^, meanwhile, are optimized for single-cell resolution and do not translate efficiently to bulk-level analyses. We developed Celestial to bridge this gap, assigning bulk-detected genes to specific cell-type origins and, where possible, attributing their age-related abundance changes to one of three mechanistically distinct sources. Due to additional consideration of population size and inter-population transcriptional similarity, Celestial outperformed Seurat-derived markers in the bulk context, particularly for small and transcriptionally similar populations such as NK and NKT cells or B and plasma cells, where subtle signature differences are easily lost in aggregate measurements. Importantly, we also showed that Celestial’s cell-type marker identification generalized to additional tissues, including defining well-characterized markers of the murine kidney. This framing recontextualizes bulk multi-omic data: rather than treating each protein or transcript as an isolated measurement, Celestial allows us to interpret changes in the context of the cell populations producing them. We view this as a tool for structured hypothesis generation—the assignments and origin-of-change calls identify candidate mechanisms that can then be tested with targeted experiments.

The trajectory of protein–RNA correlation across the lifespan offered additional insight. Per-sample correlation remained stable through early and middle life but increased significantly at 24 and 30 months. We consider two non-mutually exclusive mechanisms that may contribute to this pattern. First, the age-associated decline in protein—but not RNA—abundances of RNA-binding proteins and splicing machinery is consistent with progressive failure of post-transcriptional buffering, a process that normally decouples protein levels from their corresponding transcripts ^14,79^. This contrasts with findings in other species and tissues, where post-transcriptional dysregulation has been reported to increase discordance between protein and RNA after 24 months^12^, and where ribosome pausing has been implicated as a contributing mechanism in aging killifish^80^. The divergent trajectory we observe in spleen suggests that immune tissues may follow a distinct aging program^81^, possibly because the dramatic compositional shifts in immune cell populations impose a dominant signal that overrides subtler regulatory uncoupling^12^. Second, expansion of specific subpopulations—such as GZMK+ CD8+ T cells and C1q+ macrophages—with concurrent loss of others, including Gzma+ NK cells, naive T cells, drives coordinated changes in bulk RNA and protein for the marker genes of those populations, increasing their correlation^24,82^. Celestial’s origin-of-change analysis supports this interpretation, showing that the most prominent age-related transcript changes in cell-type-specific genes are driven by shifts in cell number rather than per-cell expression. Notably, loss of Gzma protein abundance with age—and inferred loss of NK cells—was consistent with recent human splenic proteomics data^10^. These findings highlight a general principle for aging studies: in tissues undergoing substantial cellular remodeling, changes in bulk molecular measurements may be dominated by compositional effects, and failure to account for this has the potential to misattribute regulatory mechanisms.

Sex differences emerged as a major axis of variation, shaping both the magnitude and the nature of age-related changes. In males, coordinated age-associated increases in platelet-related proteins—ALOX12, CXCL5, and the Glycoprotein Ib-IX-V-Filamin-A complex—along with complement and inflammatory proteins assigned by Celestial to macrophage and macrophage-granulocyte populations, suggest upregulation of a platelet–macrophage inflammatory axis. The strong correlation of these proteins with Cxcl5, a chemokine secreted by activated platelets^83^, alongside the absence of correlation with granulocyte markers, raises the possibility that CXCL5 preferentially recruits macrophages rather than granulocytes in this context. This male-specific pattern is consistent with the known emergence of hyperreactive platelet populations in aging males driven by distinct megakaryocyte differentiation programs^59^, and may contribute to the sex differences observed in cardiovascular and thrombo-inflammatory disease susceptibility with age. In females, we observed stronger declines in T cell-associated proteins and transcripts, particularly Wnt pathway components *Tcf7* and *Satb1*^84^, with Celestial origin-of-change analysis attributing these declines to loss of cell numbers consistent with contraction of the naive T cell compartment^19^. These sex-specific trajectories align with broader observations of sex-dimorphic immune aging^85^ and suggest that males and females may be differentially susceptible to distinct categories of age-related immune pathology—inflammatory and thrombotic in males, adaptive-immune-depleting in females.

The nonlinear inflection points we observe in the murine spleen at 21–24 months—corresponding to approximate human age of 60 years—align with molecular dysregulation recently identified by Shen et al. in longitudinal human multi-omics profiling, where the transition in humans at ∼60 years was similarly dominated by immune dysfunction (Shen et al. 2024). In our data, nonlinear changes in B cell-specific genes included decreased abundance of B cell survival factors and BCR signal transduction proteins (GO:0050853^86^; CD19, BLK, BANK1, MEF2C), several of which carry established autoimmunity risk associations^87–89^. The nonlinear age-dependent decrease in CD40 protein abundance, essential for T–B cell crosstalk, was also consistent with previous observations that the CD40⁺ B cell population contracts in elderly individuals ^90^. Together, the nonlinear erosion of B cell inhibitory receptors, survival signals, and BCR scaffold components may in part explain how immunodeficiency and autoimmune susceptibility emerge in parallel in late age^91–93^.

The case of Lef1 illustrates the value of integrating multi-omic data with cell-type assignment for generating mechanistic hypotheses. *Lef1* is among the most significantly downregulated transcripts in our dataset, yet its protein remains stable across the lifespan. Celestial assigned *Lef1* to the T cell compartment and attributed its RNA decline to loss of *Lef1*+ T cells, the majority of which lack *Ctnnb1* and *Tcf7* expression^19^. The absence of correlation between LEF1 protein and Wnt target gene RNAs is consistent with LEF1 functioning independently of canonical Wnt/β-catenin signaling in these cells^94^—possibly through HDAC activities^95^ or interactions with transcription factors such as ATF2^96^. Meanwhile, the concordant protein and RNA decline of its upstream regulators *Satb1* and *Tcf7* suggests that LEF1 protein stability may be maintained through isoform-specific protein expression, impaired proteasomal degradation, or active stabilization mechanisms^94^.

Together, these data show that the aging spleen undergoes coordinated molecular remodeling driven by overlapping changes in cell-type composition, post-transcriptional regulation, and proteostasis. By resolving the cellular origins of these changes, Celestial adds a cell-type-resolved layer to bulk multi-omic interpretation, enabling hypothesis generation about the cellular origins of molecular changes.. The sex-specific patterns and nonlinear temporal dynamics we observe reinforce the view that aging is not a uniform process of decline but a structured reorganization of tissue biology.

Looking forward, the framework established here—high temporal resolution, matched multi-omic profiling, and cell-type-resolved interpretation—is naturally extensible to a systems biology approach capturing age-related changes across multiple tissues. Specifically, cross-tissue comparisons using the same framework would clarify whether the aging trajectory we observe in spleen—particularly the increasing protein–RNA correlation—is specific to immune organs or reflects a more general principle of tissues undergoing cellular remodeling. Integration with spatial transcriptomic methods, particularly multiplexed RNA fluorescence in situ hybridization^97^, would add tissue-architectural context to the compositional changes detected by Celestial, revealing whether expanding populations such as GZMK+ CD8+ T cells and C1Q+ macrophages redistribute across splenic microanatomical compartments—for instance, from the marginal zone into the white pulp—or remain confined to their canonical niches. Our analyses identified murine aging signatures that were consistent with recent proteomics studies in aging human samples—e.g., loss of protein abundance with age for the telomerase component DKC1^10^. Thus, more broadly, extending multi-omic and cell-type-based analyses here to human cohorts would determine whether the murine spleen aging signatures translate to human immune aging.

### Limitations

Several limitations of this study should be acknowledged. Our analyses were performed in mouse spleen, and species-specific differences in cell composition or aging trajectories may limit direct translation to human biology. Our deconvolution approach integrates multiple data modalities—reference single-cell RNA sequencing, bulk transcriptomics, and bulk proteomics—each with technical and biological variability, meaning errors or assumptions can propagate through the inference pipeline. Importantly, while we focused on well annotated splenic single-cell data, our approach is inherently limited by existing cell-type annotations in the reference datasets, meaning novel or transitional populations—including those potentially emerging during aging—will be missed or misattributed. Finally, the high temporal resolution of our data offers new understanding of splenic changes, but profiling a single tissue at discrete time points limits our ability to capture the systems-level temporal dynamics of age-related remodeling. Thus, for systemic assessment of age-induced changes in murine proteome and transcriptomes, we would need to extend the current work to profile additional tissues, e.g., cell-type-specific immune infiltration in adipose tissue.

## Acknowledgements

We would like to thank members of the MacCoss, Churchill, and Schweppe labs for constructive feedback and technical assistance in assembling this work. We thank Drs. Ryan Friedman and Cole Trapnell for technical advice in developing and implementing Celestial. We would like to acknowledge the following sources of support: R35GM150919 (DKS), the W.M. Keck Foundation (DKS), an Andy Hill CARE Distinguished Researcher Award (DKS), a Cancer Consortium New Investigator Award (funded in part through P30 CA015704, DKS), the Pew Charitable Trusts (DKS), and the Jackson Laboratory Nathan Shock Center (P30AG038070, G.A.C.). We gratefully acknowledge the contribution of the RNA-seq Service at The Jackson Laboratory for expert assistance with the work described in this publication.

