## Supplementary information for "Multimodal analysis of molecular remodeling in aging spleen identified global and cell type specific changes"

**Supplementary Tables and Figures**

**Supplementary Tables**

**Supplementary Table 1.** Protein and RNA datasets including, abundance, annotations, post hoc linear and nonlinear age- and sex-effects, protein-RNA correlation and discordance measures.

**Supplementary Table 2.** GSEA analyses for pathway enrichment.

**Supplementary Table 3.** Analysis of protein complexes changes.

**Supplementary Table 4.** Celestial-assigned markers in murine spleens based on Tabula Muris Senis data.

**Supplementary Figures**

**Supplementary Figure 1.** Data quality, sex effects, and age statistics.

**Supplementary Figure 2.** Protein and RNA abundance profiles, age prediction, and correlation dynamics.

**Supplementary Figure 3.** Pathway enrichment and protein complex supplemental results.

**Supplementary Figure 4.** Celestial framework details.

**Supplementary Figure 5.** Celestial cell type-specific aging patterns.

**Supplementary Figure 6.** Detailed origin-of-change metrics across different levels.

**Materials and Methods**

#### Figure S1

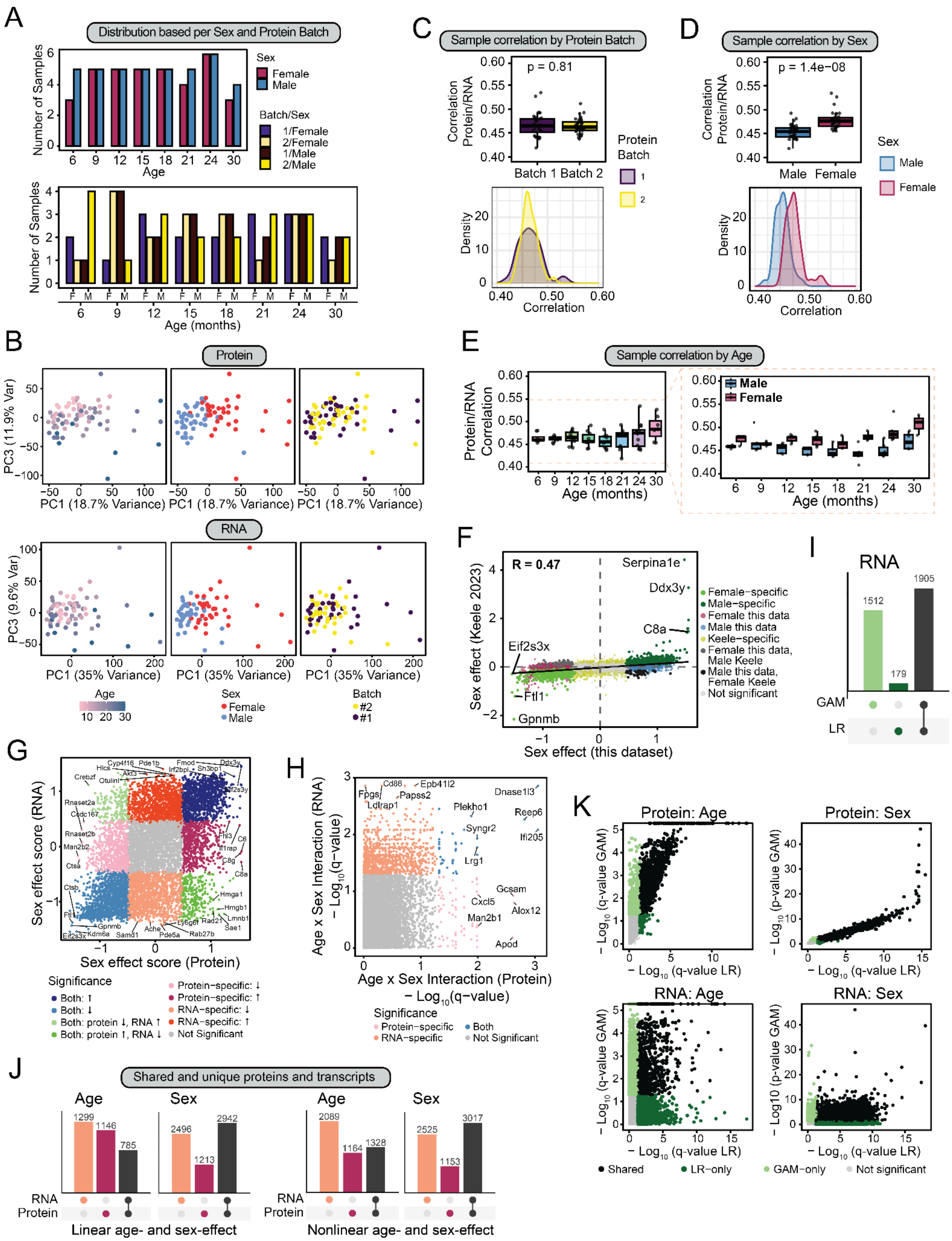

A
Distribution based per Sex and Protein Batch
Number of Samples
Age
Sex
Female
Male
Batch/Sex
1/Female
2/Female
1/Male
2/Male
Age (months)
B
Protein
PC3 (11.9% Var)
PC1 (18.7% Variance)
RNA
PC3 (9.6% Var)
PC1 (35% Variance)
Age
Sex
Female
Male
Batch
#2
#1
C
Sample correlation by Protein Batch
Correlation Protein/RNA
p = 0.81
Batch 1
Batch 2
Protein Batch
Density
Correlation
D
Sample correlation by Sex
p = 1.4e−08
Sex
Male
Female
E
Sample correlation by Age
Protein/RNA Correlation
Age (months)
F
R = 0.47
Serpina1e
Ddx3y
C8a
Eif2s3x
Ftl1
Gpnmb
Sex effect (Keele 2023)
Sex effect (this dataset)
Female−specific
Male−specific
Female this data
Male this data
Keele−specific
Female this data, Male Keele
Male this data, Female Keele
Not significant
I
RNA
GAM
LR
G
Sex effect score (RNA)
Sex effect score (Protein)
Significance
Both: ↑
Both: ↓
Both: protein ↓, RNA ↑
Both: protein ↑, RNA ↓
Protein−specific: ↓
Protein−specific: ↑
RNA−specific: ↓
RNA−specific: ↑
Not Significant
H
Age x Sex Interaction (RNA) − Log10(q−value)
Age x Sex Interaction (Protein) − Log10(q−value)
Significance
Protein−specific
RNA−specific
Both
Not Significant
K
Protein: Age
Protein: Sex
RNA: Age
RNA: Sex
− Log10 (q−value GAM)
− Log10 (p−value GAM)
− Log10 (q−value LR)
Shared
LR−only
GAM−only
Not significant
J
Shared and unique proteins and transcripts
Age
Sex
RNA
Protein
Linear age- and sex-effect
Nonlinear age- and sex-effect

**Figure S1, related to Figure 1. Data quality, sex effects, and age statistics.**

**A)** Sample overview showing sample distribution per age, sex, and batch (proteomics) across the eight timepoints.

**B)** Principal component analysis (PCA) of proteomics and transcriptomics datasets. PC1 is driven by sex in both modalities; PC3 captures age effect. Samples are colored by age, sex and batch.

**C)** Per sample Protein–RNA correlations, stratified by batch. No batch-dependent differences are observed (p-value > 0.05).

**D)** Per-sample protein–RNA correlations, stratified by sex. Female samples show higher protein–RNA correlation than males (p-value < 0.05).

**E)** Per-sample protein–RNA correlations across all eight ages, stratified by sex. Mean and standard deviation are shown per age. Correlations at 24 and 30 months are significantly higher than at ages 6–18 months. Ages 6-21: mean correlation 0.308-0.346, standard deviation (SD) 0.064-0.169; ages 24 and 30: mean correlation 0.437 and 0.484, SD 0.100 and 0.103; two-way ANOVA, BH-adjusted p < 0.05.

**F)** Scatter plot comparing sex-effect scores (LR coefficients) between this study and Keele et al. 2023 spleen data. $r_{Spearman}$=0.47. Selected genes are labeled.

**G)** Scatter plot of protein versus RNA sex-effect scores with highlighted genes having significant changes on protein and RNA levels, as well as protein-specific and RNA-specific changes between sexes.

**H)** Scatter plot of protein versus RNA negative log10 of sex-by-age interaction effects (105 significant proteins, LR) and RNA level (990 significant RNAs, LR).

**I)** Upset plots showing overlap of RNAs significantly changed with age by LR and GAM, with q-value < 0.05.

**J)** Upset plots showing overlap of significantly changing genes with age or between sexes, at the protein and RNA levels determined by LR (left) and GAM (right, q-value < 0.05).

**K)** Comparison of negative log10 of q-values calculated by LR and GAM methods per protein and per RNA.

#### Figure S2

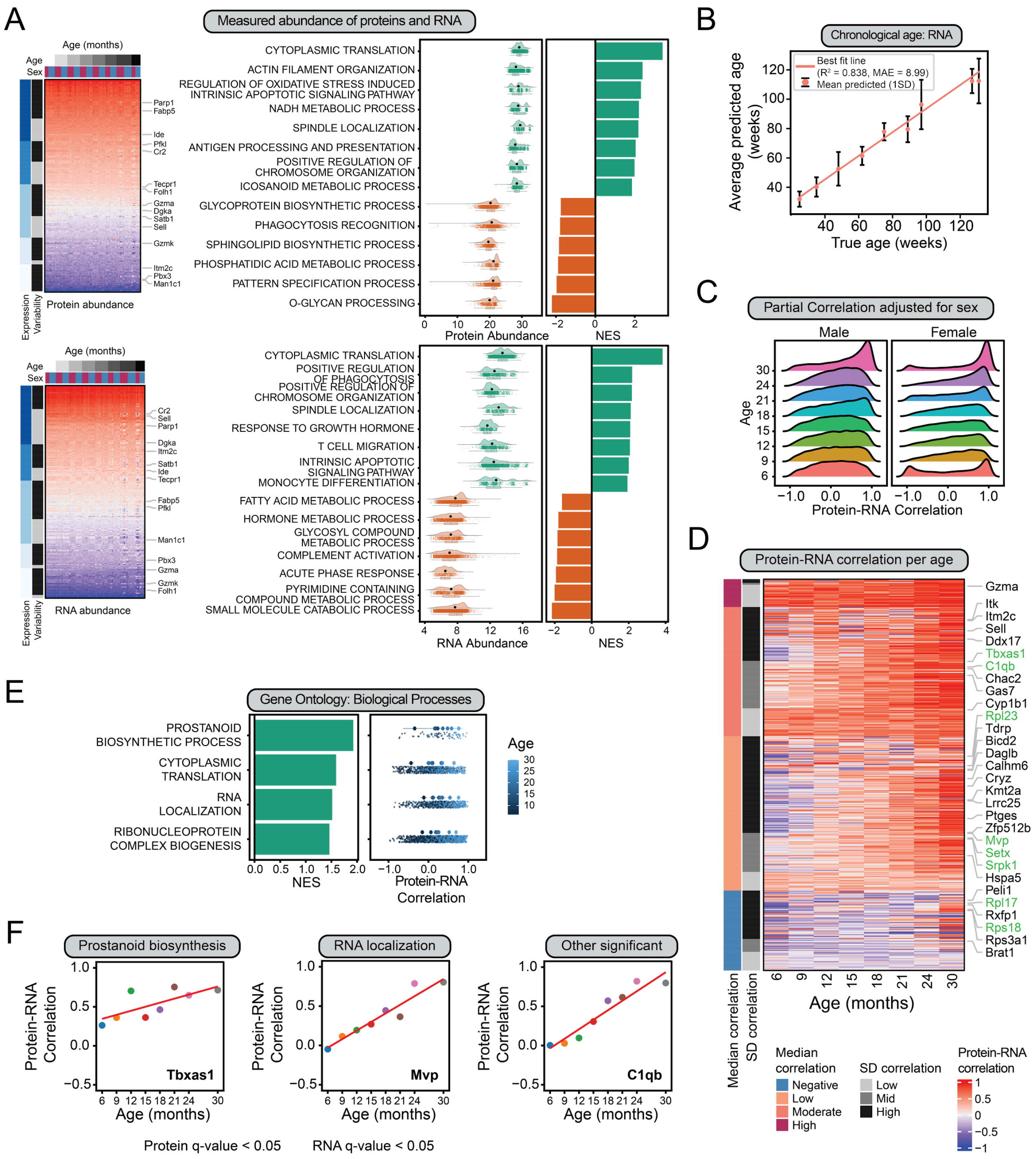

**Figure S2, related to Figure 2. Protein and RNA abundance profiles, age prediction, and correlation dynamics.**

**A)** Heatmaps of abundances of protein and RNA across all genes in our combined dataset, and the GSEA results for the abundance values. NES and distribution of the abundance of proteins or RNAs in the leading edge are shown for selected GO biological pathways.

**B)** Chronological age prediction from RNA data using elastic-net linear regression. Scatter plots show predicted versus actual age. Model performance: RNA $R^2 = 0.838$, MAE = 8.99 months.

**C)** Distribution of per-gene partial correlations between protein and RNA abundance adjusted for sex, calculated separately at each age and stratified by sex.

**D)** Heatmap showing age-dependent protein–RNA partial correlations adjusted for sex, with highlighted genes with specific age-specific patterns across the lifespan.

**E)** GSEA results for slope (LR) values of protein–RNA correlation change with age. NES shows enriched gene sets with increasing protein–RNA correlation. Partial correlation per age of the genes in the leading age shows age-specific change.

**F)** Examples of age-dependent protein–RNA partial correlations change with age of gene in prostanoid pathway (Tbxas1), RNA localisation (Mvp) and other gene with high slope (C1qb).

### Figure S3

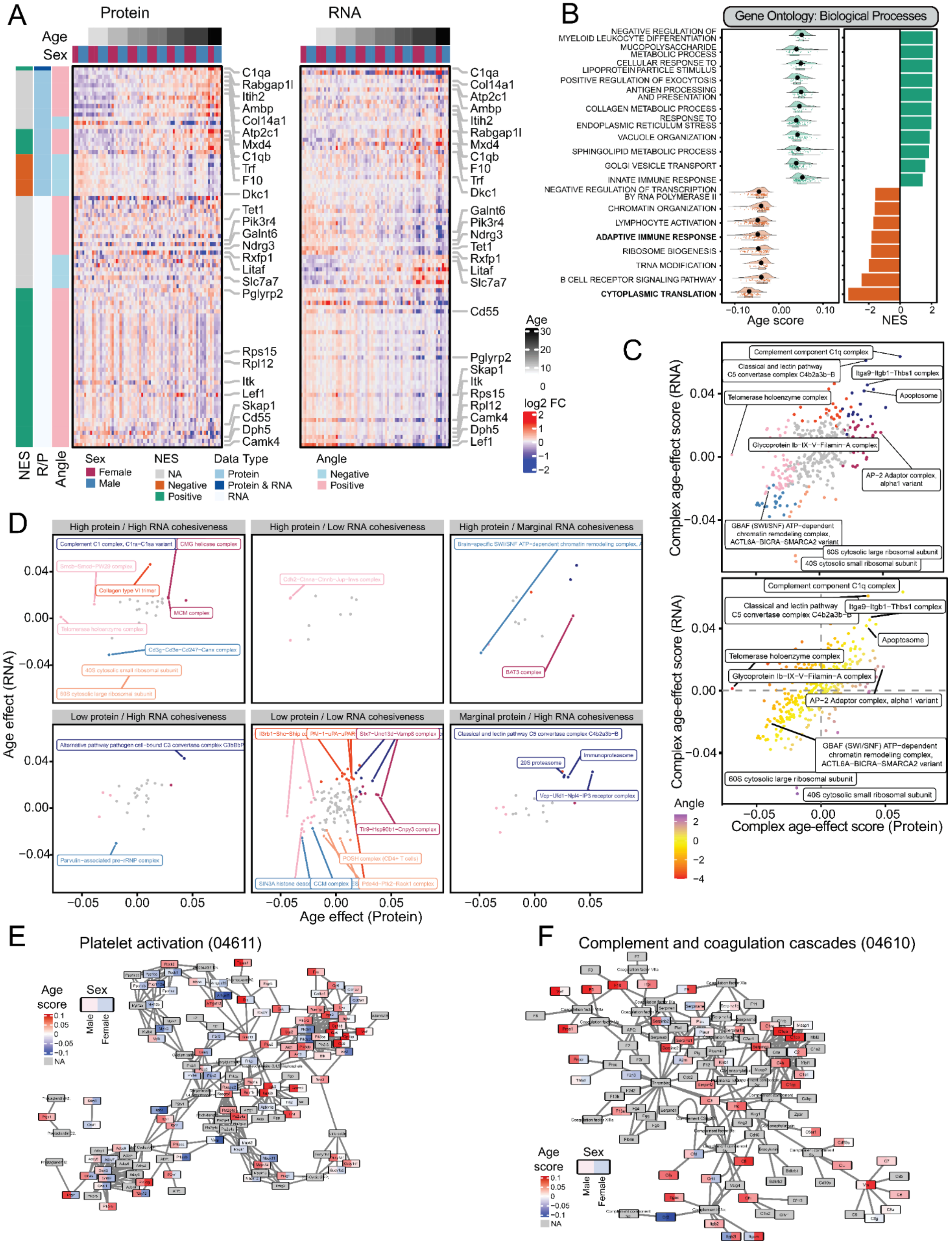

A
Protein
RNA
Age
Sex
C1qa
Rabgap1l
Itih2
Ambp
Col14a1
Atp2c1
Mxd4
C1qb
Trf
F10
Dkc1
Tet1
Pik3r4
Galnt6
Ndrg3
Rxfp1
Litaf
Slc7a7
Pglyrp2
Cd55
Rps15
Rpl12
Itk
Lef1
Skap1
Camk4
Dph5
NES
R/P
Angle
Sex
Female
Male
NES
NA
Negative
Positive
Data Type
Protein
Protein & RNA
RNA
Angle
Negative
Positive
log2 FC
B
Gene Ontology: Biological Processes
NEGATIVE REGULATION OF MYELOID LEUKOCYTE DIFFERENTIATION
MUCOPOLYSACCHARIDE METABOLIC PROCESS
CELLULAR RESPONSE TO LIPOPROTEIN PARTICLE STIMULUS
POSITIVE REGULATION OF EXOCYTOSIS
ANTIGEN PROCESSING AND PRESENTATION
COLLAGEN METABOLIC PROCESS
RESPONSE TO ENDOPLASMIC RETICULUM STRESS
VACUOLE ORGANIZATION
SPHINGOLIPID METABOLIC PROCESS
GOLGI VESICLE TRANSPORT
INNATE IMMUNE RESPONSE
NEGATIVE REGULATION OF TRANSCRIPTION BY RNA POLYMERASE II
CHROMATIN ORGANIZATION
LYMPHOCYTE ACTIVATION
ADAPTIVE IMMUNE RESPONSE
RIBOSOME BIOGENESIS
TRNA MODIFICATION
B CELL RECEPTOR SIGNALING PATHWAY
CYTOPLASMIC TRANSLATION
Age score
NES
C
Complex age-effect score (RNA)
Complex age-effect score (Protein)
Complement component C1q complex
Classical and lectin pathway C5 convertase complex C4b2a3b-B
Itga9-Itgb1-Thbs1 complex
Apoptosome
Telomerase holoenzyme complex
Glycoprotein Ib-IX-V-Filamin-A complex
AP-2 Adaptor complex, alpha1 variant
GBAF (SWI/SNF) ATP-dependent chromatin remodeling complex, ACTL6A-BICRA-SMARCA2 variant
60S cytosolic large ribosomal subunit
40S cytosolic small ribosomal subunit
Angle
D
High protein / High RNA cohesiveness
High protein / Low RNA cohesiveness
High protein / Marginal RNA cohesiveness
Low protein / High RNA cohesiveness
Low protein / Low RNA cohesiveness
Marginal protein / High RNA cohesiveness
Complement C1 complex, C1ra-C1sa variant
CMG helicase complex
Collagen type VI trimer
MCM complex
Telomerase holoenzyme complex
Cd3g-Cd3e-Cd247-Canx complex
40S cytosolic small ribosomal subunit
60S cytosolic large ribosomal subunit
BAT3 complex
Alternative pathway pathogen cell-bound C3 convertase complex C3bBbP
Parvulin-associated pre-rRNP complex
Tlr9-Hsp90b1-Cnpy3 complex
POSH complex (CD4+ T cells)
SIN3A histone deac
CCM complex
Pde4d-Ptk2-Rock1 complex
Classical and lectin pathway C5 convertase complex C4b2a3b-B
20S proteasome
Immunoproteasome
Vcp-Ufd1-Npl4-IP3 receptor complex
Age effect (RNA)
Age effect (Protein)
E
Platelet activation (04611)
Age score
Sex
Male
Female
NA
F
Complement and coagulation cascades (04610)

**Figure S3, related to Figure 3. Pathway enrichment and protein complex supplemental results.**

**A)** Heatmap of genes with high and low protein–RNA discordance, belonging to specific pathways (indicated by NES values), or not matched to any biological pathway.

**B)** GSEA results for RNA age-effect scores, showing all significantly enriched GO biological processes. Shown are NES values for selected GO biological pathways and distribution of age-effect score values for genes in the leading edge.

**C)** Scatter plot of protein versus RNA age-effect scores per protein complex (top), and discordance between their age-effect scores. Highlighted are protein complexes having significant changes on protein and RNA levels, as well as protein-specific and RNA-specific changes.

**D)** Age-effect scores at protein and RNA-level, stratified by cohesiveness values.

**E-F)** Network plots of the “Platelet activation” and “Complement and coagulation cascade” pathways. Genes are colored by protein age-effect score per sex.

### Figure S4

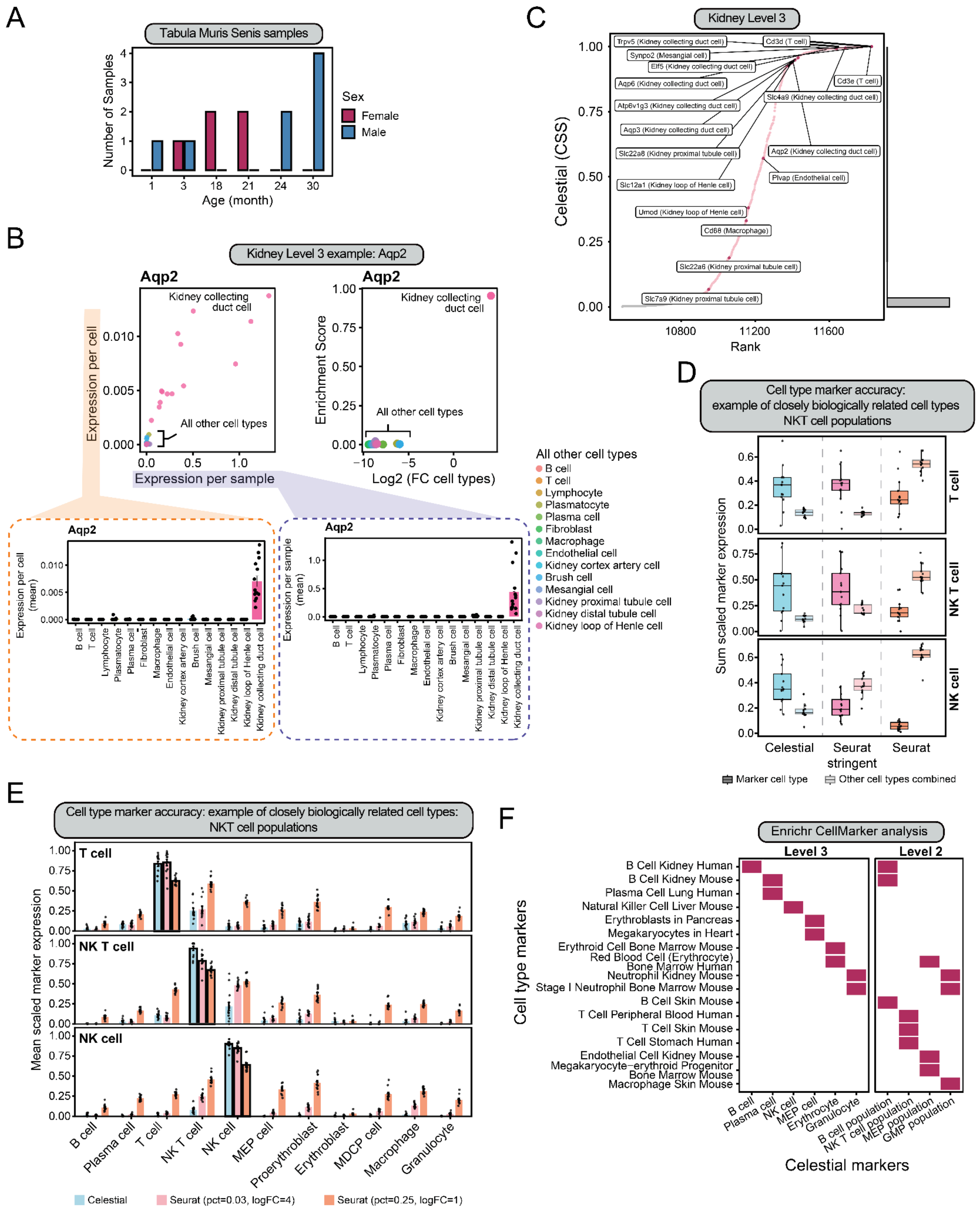

A
Tabula Muris Senis samples
Number of Samples
Sex
Female
Male
Age (month)
B
Kidney Level 3 example: Aqp2
Aqp2
Kidney collecting duct cell
All other cell types
Expression per cell
Expression per sample
Enrichment Score
Log2 (FC cell types)
Expression per cell (mean)
Expression per sample (mean)
All other cell types
B cell
T cell
Lymphocyte
Plasmatocyte
Plasma cell
Fibroblast
Macrophage
Endothelial cell
Kidney cortex artery cell
Brush cell
Mesangial cell
Kidney proximal tubule cell
Kidney distal tubule cell
Kidney loop of Henle cell
Kidney collecting duct cell
C
Kidney Level 3
Celestial (CSS)
Rank
Trpv5 (Kidney collecting duct cell)
Cd3d (T cell)
Synpo2 (Mesangial cell)
Elf5 (Kidney collecting duct cell)
Aqp6 (Kidney collecting duct cell)
Cd3e (T cell)
Atp6v1g3 (Kidney collecting duct cell)
Slc4a9 (Kidney collecting duct cell)
Aqp3 (Kidney collecting duct cell)
Slc22a8 (Kidney proximal tubule cell)
Aqp2 (Kidney collecting duct cell)
Plvap (Endothelial cell)
Slc12a1 (Kidney loop of Henle cell)
Umod (Kidney loop of Henle cell)
Cd68 (Macrophage)
Slc22a6 (Kidney proximal tubule cell)
Slc7a9 (Kidney proximal tubule cell)
D
Cell type marker accuracy: example of closely biologically related cell types NKT cell populations
Sum scaled marker expression
T cell
NK T cell
NK cell
Celestial
Seurat stringent
Seurat
Marker cell type
Other cell types combined
E
Cell type marker accuracy: example of closely biologically related cell types: NKT cell populations
Mean scaled marker expression
T cell
NK T cell
NK cell
B cell
Plasma cell
MEP cell
Proerythroblast
Erythroblast
MDCP cell
Macrophage
Granulocyte
Celestial
Seurat (pct=0.03, logFC=4)
Seurat (pct=0.25, logFC=1)
F
Enrichr CellMarker analysis
Level 3
Level 2
Cell type markers
B Cell Kidney Human
B Cell Kidney Mouse
Plasma Cell Lung Human
Natural Killer Cell Liver Mouse
Erythroblasts in Pancreas
Megakaryocytes in Heart
Erythroid Cell Bone Marrow Mouse
Red Blood Cell (Erythrocyte) Bone Marrow Human
Neutrophil Kidney Mouse
Stage I Neutrophil Bone Marrow Mouse
B Cell Skin Mouse
T Cell Peripheral Blood Human
T Cell Skin Mouse
T Cell Stomach Human
Endothelial Cell Kidney Mouse
Megakaryocyte−erythroid Progenitor Bone Marrow Mouse
Macrophage Skin Mouse
B cell
Plasma cell
NK cell
MEP cell
Erythrocyte
Granulocyte
B cell population
NK T cell population
MEP population
GMP population
Celestial markers

**Figure S4, related to Figure 4. Celestial framework details.**

**A)** Sample overview from TMS spleen data, showing sample distribution per age and sex across aging.

**B)** Plots showing features distributions (including detailed view) used by Celestial for cell-type-specific gene classification. Shown are plots for *Aqp2* from kidney TMS dataset, indicating specificity of *Aqp2* expression in kidney collecting duct cells.

**C)** Example of Celestial applied to kidney TMS dataset, showing Celestial plot of random forest scores versus rank of assignment genes at Level-3 (cell type-specific genes).

**D)** Plot corresponds to bulk-marker accuracy, showing sum of all genes per cell type of the marker or in all other cell types. Each marker was scaled between 0 and 1. For accurate marker calling, higher expression in the marker cell-type (darker colors) than other cell-types (lighter colors) is expected.

**E)** Plot corresponds to cell type-marker accuracy, showing marker specificity across cell types per donor for all three methods. Each marker was scaled between 0 and 1, and the mean of all markers was used to plot.

**F)** Example Celestial plots wherein ranked gene-by-cell-type relationships are indexed based on Celestial cell-type specificity score (CSS) for Level-3 (n=584 cell type-specific genes, top) and Level-0 (n=1638 global genes, bottom). Statistical significance was assessed by fitting a beta distribution to background scores from genes not meeting the specificity criteria and computing FDR-adjusted p-values.

Figure S5

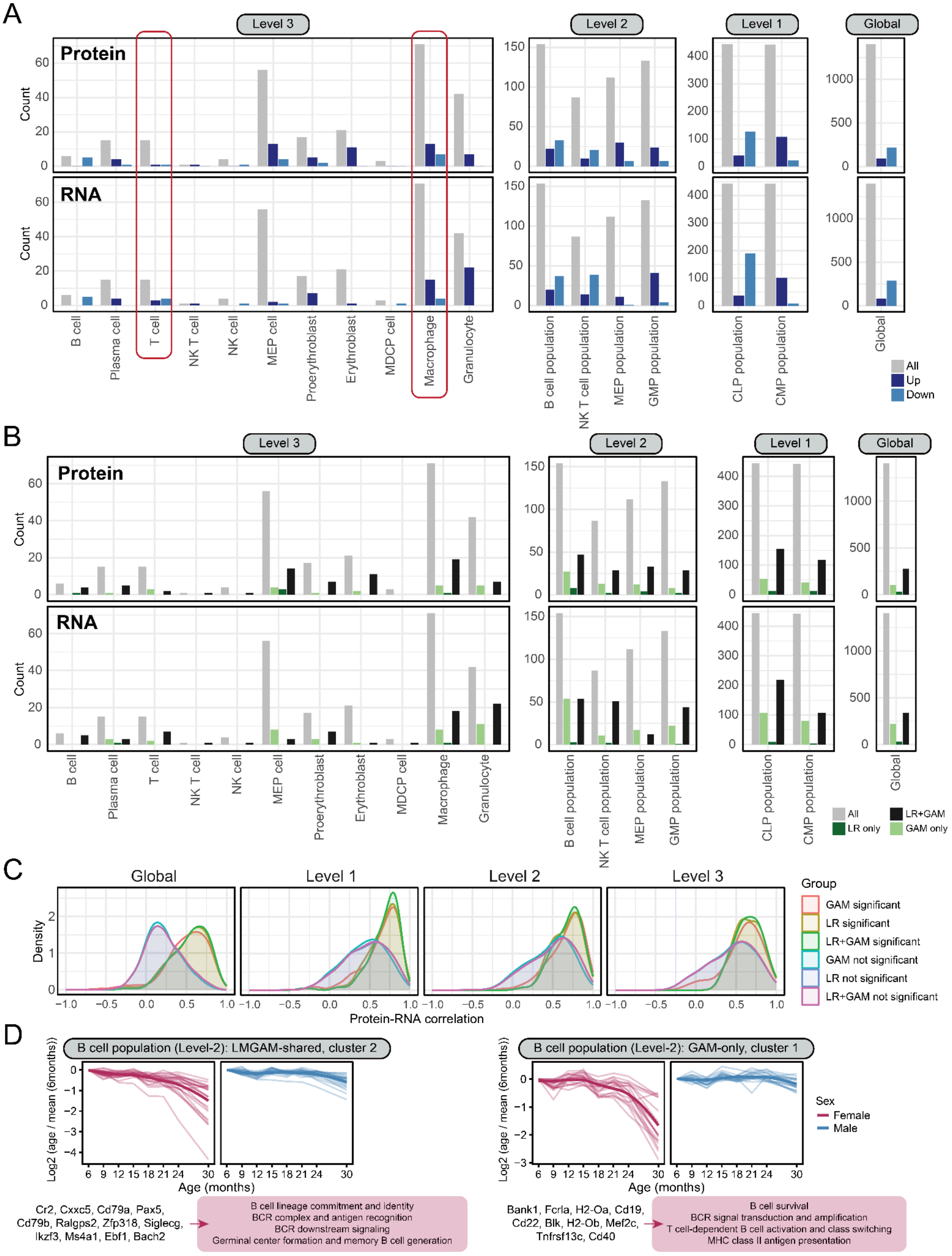
A
Level 3
Level 2
Level 1
Global
Protein
RNA
Count
B cell
Plasma cell
T cell
NK T cell
NK cell
MEP cell
Proerythroblast
Erythroblast
MDCP cell
Macrophage
Granulocyte
B cell population
NK T cell population
MEP population
GMP population
CLP population
CMP population
All
Up
Down
B
All
LR only
LR+GAM
GAM only
C
Density
Protein-RNA correlation
Group
GAM significant
LR significant
LR+GAM significant
GAM not significant
LR not significant
LR+GAM not significant
D
B cell population (Level-2): LMGAM-shared, cluster 2
B cell population (Level-2): GAM-only, cluster 1
Log2 (age / mean (6months))
Age (months)
Sex
Female
Male
Cr2, Cxxc5, Cd79a, Pax5, Cd79b, Ralgps2, Zfp318, Siglecg, Ikzf3, Ms4a1, Ebf1, Bach2
B cell lineage commitment and identity
BCR complex and antigen recognition
BCR downstream signaling
Germinal center formation and memory B cell generation
Bank1, Fcrla, H2-Oa, Cd19, Cd22, Blk, H2-Ob, Mef2c, Tnfrsf13c, Cd40
B cell survival
BCR signal transduction and amplification
T cell-dependent B cell activation and class switching
MHC class II antigen presentation

**Figure S5, related to Figure 5. Celestial cell type-specific aging patterns.**

**A)** Plots showing numbers of Celestial-identified genes at each level (Level-3 through Level-0), including all genes, as well as numbers of significantly upregulated and downregulated (determined by LR).

**B)** Plots showing numbers of Celestial-identified genes at each level (Level-3 through Level-0), including all genes and those significantly changing based on LR (linear), GAM (non-linear), or Both.

**C)** Density plots of protein–RNA partial correlations across all levels (Level-3 through Level-0), showing difference in distribution of genes significantly changing in both proteomics and transcriptomics data (shared) identified by LR, GAM, or shared between LR and GAM, and the rest of genes present in the corresponding level.

**D)** Protein abundance trajectories across age for subset of markers assigned to B cell population (Level-2): subset of significantly changing proteins using LR and GAM in cluster 2, or GAM only in cluster 1 (**Fig. 1F**). Proteins from these clusters were 1) significantly enriched in the B cell-related signaling, activation, differentiation and proliferation processes identified with Enrichr (indicated in the text boxes)[66], and 2) contribute to additional processes manually assigned from literature.

**Figure S6**

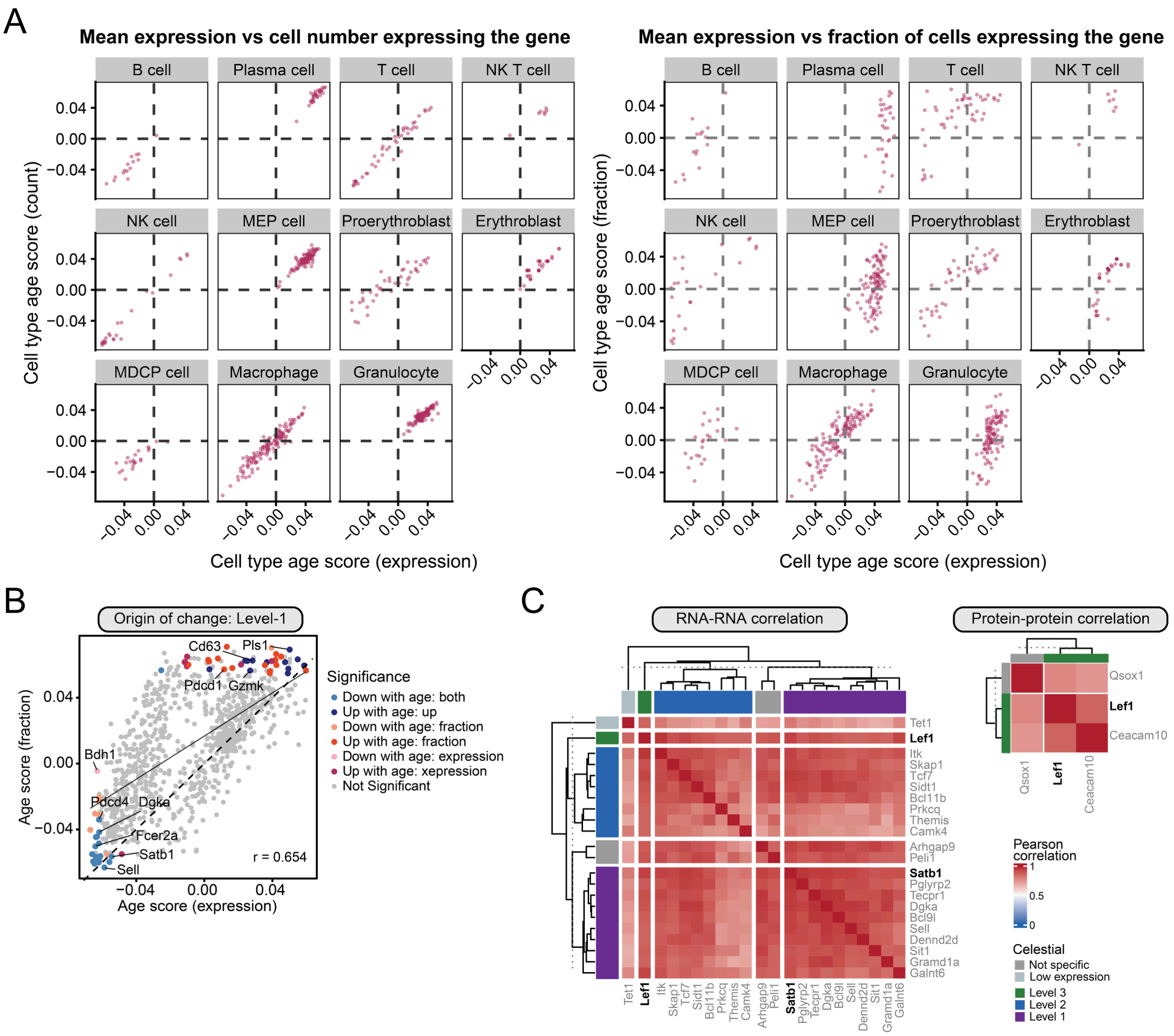


**Figure S6, related to Figure 6. Detailed origin-of-change metrics across different levels.**

**A)** Scatter plots of age-effect scores across cell type-specific genes for all three origin-of-change metrics, showing how correlated are feature metrics per each cell type (Level-3).

**B)** Origin-of-change metrics across age for Satb1 at the Level-1 (lymphoid population CLP) level, showing decline across all three metrics consistent with reduction of Satb1+ T cells.

**C)** RNA–RNA and protein-protein correlation matrices (adjusted for sex) for genes with the highest transcriptomic correlation to Lef1, including Tcf7, Satb1, Tet1, and Itk, and their cell type assignments. The protein correlation matrix demonstrated that the strong RNA co-regulation pattern is not reflected at the protein level.

#### Materials and methods

##### Aging mouse models used in this study

Male and female C57BL/6J mice ages 6, 9, 12, 15, 18, 21, 24 and 30 months were obtained from The Jackson Laboratory. A total of 76 mice were used in the study, with on average five animals per sex per age. Animals were housed on pine shavings in a pathogen free room at 21°C with a 12-hour light/dark cycle (6am to 6pm). Food (LabDiet 5KOG) and acidified water were available ad libitum. Animals were euthanized by cervical dislocation and whole spleens were collected from each animal. Each spleen was flash-frozen in liquid nitrogen and cryopulverized. All animal experiments were in accordance with the National Institutes of Health Guide for the Care and Use of Laboratory Animals (National Research Council). All the protocols and procedures were reviewed and approved by the Animal Care and Use Committee at The Jackson Laboratory.

##### Sample preparation for proteomic analysis

Cryopulverized spleen tissue aliquots designated for proteomics analysis were resuspended in lysis buffer (2% SDS, 100 mM Tris, 1X HALT protease/phosphatase inhibitors), vortexed, and briefly probe sonicated. Protein concentration was measured using a BCA assay. Homogenate (100 µg of protein) was spiked with 1600 ng yeast enolase (process control), reduced with 10 mM TCEP, alkylated with 15 mM IAA, and quenched with 10 mM DTT. Proteins were aggregated on MagResyn Hydroxyl beads (Resyn Biosciences) with 70% acetonitrile, washed, and trypsin (ratio 1:10) digested to peptides, using a Thermo KingFisher Apex.

##### Data-independent acquisition mass spectrometry

Each sample injection contained 3 µg total protein (48 ng enolase process control) and 450 fmol Peptide Retention Time Calibrant mixture (PRTC, ThermoFisher). Samples were analyzed on a Thermo Vanquish Neo UHPLC System using a 15 cm × 150 µm PepSep (Bruker) column and PepMap (ThermoFisher) Neo 300 µm x 5 mm of 5 µm C18 trap with a LOTUS nESI 20 µm emitter (Fossil). Mobile phases were 0.1% formic acid (A) and 0.1% formic acid in 80% acetonitrile (B). System suitability was monitored with four QC runs before sample analysis and after every six samples. Peptides were eluted from the column with a 50°C heated source and electrosprayed with 3 kV voltage into a Thermo Orbitrap Fusion Lumos mass spectrometer. Data acquisition was controlled by Xcalibur 3.1.2412.24. For chromatogram library generation, six pooled sample injections were analyzed using 100 m/z windows (400-1000 *m/z*) with data-independent acquisition (DIA): 30,000 resolution full-scan MS1, followed by data-independent 25 MS/MS scans at 30,000 resolution (4 *m/z* overlapping isolation windows, automatic max injection time, 27% normalized collision energy). Individual samples were analyzed using DIA with 350-2000 *m/z* full-scan MS1 at 30,000 resolution, followed by data-independent 50 MS/MS scans at 30,000 resolution (8 *m/z* overlapping isolation windows, automatic max injection time, 27% normalized collision energy).

##### Proteomics data processing

Thermo raw files were demultiplexed [98] and converted to mzML files using ProteoWizard's msConvert [99] version 3.0.23187 using the following arguments: --mzML --zlib --ignoreUnknownInstrumentError --filter "peakPicking true 1-" --64 --filter "demultiplex optimization=overlap_only" –simAsSpectra. A Nextflow workflow (version 23.04.4) was used to build an instrument-specific, optimized, spectral library in dlib format generated using the UniProt reviewed mouse proteome (UP000000589_10090, Oct 2024) plus DIA-NN [100] version 1.8.1 and Carafe version 0.0.1 as previously described [101]. The pooled gas phase fractionated chromatogram library was created using EncyclopeDIA [102] (version 2.12.30) with default settings using the Carafe generated spectral library based on the UniProt mouse proteome FASTA. This chromatogram library was used to quantify peptides using EncycopeDIA analysis of the wide window DIA runs from the individual sample runs. Extracted ion chromatograms and peak areas were exported using Skyline-daily [103] (64-bit version 23.1.1.537). The conversion of Skyline reports to unnormalized and normalized precursor and protein level reports was done using an inhouse produced Nextflow workflow available at: https://github.com/uw-maccosslab/nf-dia-batch-correction.

**Proteomic Data Availability**

Raw MS data, sample metadata, Skyline documents, quantified precursor areas and summed protein areas are available at Panorama Public [104105] Access URL: https://panoramaweb.org/mouse-aging-spleen.url.

**Sample preparation for RNA sequencing analysis**

Cryopulverized spleen tissue aliquots designated for transcriptomics were processed at the Genome Technologies service (Jackson Laboratory). RNA extraction was performed using the MagMAX mirVana Total RNA Isolation Kit (ThermoFisher Scientific) with automated purification on the KingFisher Flex system. Tissue samples were lysed in TRIzol, the aqueous phase was separated with chloroform, and RNA was purified from the aqueous layer using magnetic bead-based isolation following the manufacturer's protocol. RNA yield and integrity were assessed by Nanodrop 8000 spectrophotometer and Agilent Total RNA Nano assay.

Libraries were prepared using the KAPA mRNA HyperPrep Kit (Roche). Poly-A enriched mRNA was captured from total RNA using oligo-dT magnetic beads, then subjected to fragmentation, synthesis of first- and second-strand cDNA, ligation of Illumina-specific adapters containing unique dual index barcodes, and PCR amplification. Library integrity was assessed using D5000 ScreenTape (Agilent Technologies) and concentration measured using Qubit dsDNA HS Assay (Thermo Fisher Scientific). Pooled libraries were sequenced on the Illumina NovaSeq 6000 platform with S1 Reagent Kit v1, generating 20-40M (target 30M) 1 × 100bp single-end reads per sample.

**Transcriptomics data processing**

Gene expression quantification used the Expectation-Maximization algorithm for Allele Specific Expression (EMASE)[106], to obtain gene-level read counts from the RNA-seq data.

Reads were aligned to the transcriptome derived from mouse reference genome GRCm38. Transcripts were removed from further analysis if they did not have at least XX reads and gene-level counts were normalized relative to total read counts using the variance stabilizing transform (VST) as implemented in DESeq2[107]. For most analyses, to minimize the impact of outliers, we transformed the VST normalized data to rank normal scores[108].

#### Integration of proteomics and transcriptomics data

Spleen proteomic and transcriptomic datasets were processed and integrated at the gene level. Data were filtered for genes that were uniquely mapped and quantified in all samples. Datasets were aligned using Ensembl gene ID and UniProt ID mapping, resulting in 8,528 genes for multi-omics analysis. Data were either rank-z normalized or $\log_2$-transformed and median-normalized per gene. The processed data is viewable through the shiny app available online at: https://kvlajic.shinyapps.io/spleen-aging-atlas/

Principal component analysis (PCA) was performed on both protein and RNA $\log_2$-transformed and median-normalized datasets to assess overall sample structure and identify potential batch effects. Sample pairwise Pearson correlations were calculated within each modality (protein-protein and RNA-RNA) and across modalities (protein–RNA) using non-transformed abundance values. These per-sample correlations were used to evaluate data quality and the relationship between protein and RNA levels across samples.

#### Chronological age estimation

Chronological age was estimated from protein and RNA abundance data using machine learning approaches. Predictive models were built using elastic net linear regression to estimate chronological age from the selected molecular features based on protein and RNA abundance.

#### Statistical modeling of age- and sex-associated molecular changes

To identify proteins and transcripts that change with age and sex, linear regression (LR) and generalized additive models (GAMs) [109] were fit for each gene. Data were rank-z normalized per gene. For LR, we tested age effects, sex effects, and age-by-sex interactions using four models for each gene:

- Model 1: $Yi = \beta_0 + \beta_{Age} \times Age_i + \varepsilon_i$
- Model 2: $Yi = \beta_0 + \beta_{Sex} \times Sex_i + \varepsilon_i$
- Model 3: $Yi = \beta_0 + \beta_{Sex} \times Sex_i + \beta_{Age} \times Age_i + \varepsilon_i$
- Model 4: $Yi = \beta_0 + \beta_{Sex} \times Sex_i + \beta_{Age} \times Age_i + \beta_{Sex \times Age}(Sex \times Age)_i + \varepsilon_i$

Y represents rank-z normalized expression, Age is a continuous variable (in months), and Sex is a categorical variable. Significance of effects on abundances were tested using anova F-test. Significance of age effects were tested by comparing Model 3 to Model 2, sex effects by comparing Model 3 to Model 1, age-by-sex interaction effects by comparing Model 4 to Model 3. Age- and sex-effects were extracted from regression coefficients from Model 3. Sex-specific age-effects were estimated using estimated marginal trends from Model 4. Standardized coefficients were calculated using a regression coefficient divided by standard error.

For GAMs, non-linear age trajectories were estimated using smooth functions:

- Model 5: Y ~ Sex + s(Age, k = 5, m = 3) + s(Age, by = Sex, k = 5, m = 3),

where s() represents a smooth function with flexibility controlled by basis dimension k = 5 and penalty order m = 3. Models were fit using restricted maximum likelihood (REML). Age effects were estimated from the smooth term p-values, and sex-by-age interaction effects were estimated from the sex-specific smooth term.

Correction for multiple testing was performed using the Benjamini-Hochberg. Genes were classified as significantly changing based on adjusted p-value < 0.05.

**Protein–RNA concordance and discordance**

Relationship between protein and RNA dynamics during aging was analyzed at the gene level using $log_2$-normalized data. For each gene, partial correlations between protein and RNA abundance across all samples were calculated while controlling for sex using partial correlation tests. Correction for multiple testing was performed using the Benjamini-Hochberg.

To explore how protein–RNA coupling changed with age, partial correlations were calculated separately at each age. For each age, partial correlations between protein and RNA were calculated for each gene while controlling for sex. Linear models were then fitted for each gene to test whether correlations changed with age:

Model 6: Model 1: $Yi = \beta_0 + \beta_{Age} \times Age_i + \varepsilon_i$ (Partial_Correlation ~ Age)

Age coefficients, standard errors, and standardized coefficients (coefficient divided by standard error) were extracted, and p-values were adjusted for multiple comparison using the Benjamini-Hochberg.

Discordance in aging trajectories of protein and RNA was quantified for each gene by calculating the angle between protein and RNA age-effect slopes (linear regression, Model 3) using arctangent (atan2):

$$Discordance = 180/\pi \times arctan(\beta(protein) - \beta(RNA)) / (1 + \beta(protein) - \beta(RNA))$$

where β are age-effect coefficients from Model 3. Angles closer to zero indicate concordant relationship, larger angles suggest discordant aging trajectories. In this dataset, observed angles ranged ±7-8°, with a maximum constraint of ±12.5° given the range of estimated age-effect coefficients.

Statistical significance of protein–RNA discordance was assessed per gene using a Wald test for the difference between two regression coefficients:

Model 7: $Z = (\beta_{protein} - \beta_{RNA}) / \sqrt{SE^2_{protein} + SE^2_{RNA}}$

SE = regression coefficient / standardized regression coefficient

Two-sided p-values were derived from the standard normal distribution and adjusted for multiple testing using the Benjamini-Hochberg. To assess significance of the RNA-protein discordances, we used a Wald test for model parameters[110]. The Discordance metric (angle) and Wald Z-statistic correlated positively ($r_{Spearman}$=0.999), confirming that both measures capture the same relationship. Genes with q-value < 0.05 were classified as protein-dominant if protein age-effect score was greater than RNA age-effect score or RNA-dominant if RNA age-effect score was greater than protein age-effect score.

Wnt-target genes were obtained from MSigDB (Mouse Gene Set: Ctnnb1 target genes). Protein and RNA were correlated with RNA, and p-values were adjusted for multiple comparisons (in comparison to the whole data) using the Benjamini-Hochberg. RNA with q-value < 0.05 were classified as correlating with protein or RNA it was compared to.

##### Gene set enrichment analysis

Gene set enrichment analysis (GSEA) [111] was performed using the fgsea R package [112] with Gene Ontology (GO) gene sets obtained from the Molecular Signatures Database (MSigDB) for organism Mus musculus. GO gene sets were divided into three categories: biological processes (BP), molecular functions (MF), and cellular components (CC). Only gene sets with 15-500 genes (10-500 for correlation analysis) were included. Enrichment was assessed using 10,000 permutations (100,000 for correlation analysis).

GSEA was performed using four different gene ranking metrics: abundance, slope (per modality), discordance angle, and slope (protein–RNA correlation linear model). For abundance, genes were ranked by the mean rank-z normalized abundance values. For age effect slope per modality, genes were ranked by the standardized coefficients from linear Model 3. For discordance, genes were ranked by the scaled angle between protein and RNA age trajectories. For slopes modeling correlation dynamics, genes were ranked by standardized coefficients from linear Model 6. Results from Celestial were ranked by their random forest score (CSS), combining Levels 1-3 (using the highest CSS value) as positive and Level-0 as negative edges.

All rankings were performed separately for protein and RNA datasets or for values exploring their relationships, and genes were sorted in descending order. Enrichment was assessed by calculating normalized enrichment scores (NES), p-values, and adjusted p-values for each GO gene set. Gene ontologies were selected based on adjusted p-value, < 0.01 for abundance, slope, and angle, adjusted p-value < 0.05 for correlation and discordance.

##### Protein complex analysis

Protein complex annotations were obtained from CORUM and Complex Portal databases for Mus Musculus [53,54]. Complexes from both databases were matched and grouped based on their member proteins. Jaccard index was calculated for all complex pairs, and complexes with Jaccard index ≥ 0.75 were merged to remove redundancy. For merged complexes, the complex with the largest number of proteins was selected as the representative. Only complexes with at least 3 detected components in the dataset were used for the analysis.

Age effects on protein complex abundances were analyzed using linear mixed-effects models (LMMs). Protein and RNA abundance data were rank-z normalized per gene. For each complex, data were filtered for subunits of the complex. Age effects were tested using models for each complex:

Model 8 (full): $Y_i = \beta_0 + \beta_1 \times Sex_i + \beta_2 \times (Age)_{ij} + \Sigma\beta_{4k} Batch_{ijk} + b_{0j} + \varepsilon_i$

Y ~ Age + Sex + ProtBatch + (1|MouseID)

Model 9: Y ~ Sex + ProtBatch + (1|MouseID)

where Age was modeled as a continuous variable (in months), Sex and ProtBatch were fixed effects, and MouseID was included as a random effect to account for repeated measurements (all complex components per sample). Age effects were explored by comparing Model 8 to Model 9 using likelihood ratio tests. Age coefficients, standard errors, and p-values were extracted from Model 1. P-values were adjusted for multiple comparisons using the Benjamini-Hochberg. Complexes labeled as significantly changing were based on adjusted p-value < 0.05. The relationships between aging trajectories of protein and RNA components of protein complexes were analyzed using arctangent.

Complex cohesiveness was calculated as a median partial correlation between all protein components within a protein complex. Partial correlations between all protein pairs were calculated across all samples while controlling for sex using partial correlation tests. For each complex, pairwise partial correlations were extracted for all subunits of the complex, and the median correlation was calculated as the complex cohesiveness. It represents a degree of coordinated change between all subunits within the complex. Higher cohesiveness indicates greater co-regulation of complex components.

**Development of the Celestial Framework**

To distinguish cell type-specific from globally expressed genes in bulk transcriptomic and proteomic data, we developed Celestial, a computational framework that integrates single-cell RNA-seq reference data with bulk multi-omics measurements. Celestial is available at github.com/SchweppeLab/celestial

*Single cell reference data.* scRNA-seq data from the murine spleen was obtained from the Tabula Muris Senis (TMS) atlas[19]. Only droplet-based sequencing data was used to ensure consistency in library preparation and gene coverage across ages per cell type.

*Cell type-specific gene identification.* For each gene, expression data was normalized per cell type by dividing by the mean transcriptome size within that cell type to account for differences in sequencing depth. Multiple cell type specificity features were calculated across cell types: (1) total expression per cell type (sum of normalized counts), (2) expression per cell (total expression divided by cell number), (3) enrichment score (ratio of cell type expression to total expression, scaled by number of cell types), (4) $\log_2$ differential expression ($\log_2$ ratio of cell type expression to the sum of all other cell types), and (5) fraction of cells expressing the gene (proportion of cells with non-zero counts). Z-score transformations and scaled maximum differences from the top cell type were calculated per gene across all cell types. Genes passing at least 4 out of 5 specificity criteria were considered candidates for cell type-specific markers.

*Global gene identification.* Global genes were identified by testing for flat expression profiles across cell types, using features calculated for Level-3. For each gene, all cell types were evaluated for similarity using fold-change ratios between consecutive cell type ranks. Coverage was assessed as the proportion of cell types with ≥10% of cells expressing the gene, requiring ≥50% coverage. Genes passing at least 4 out of 5 criteria—expression per cell flatness, fraction expressing flatness, enrichment score flatness, $\log_2$ differential closeness, and coverage threshold—were classified as globally expressed.

*Cell-type specificity score.* Cell type-specific and global gene scores were generated using a random forest-based feature integration approach. All genes were labeled based on whether they met the specificity criteria described above, and a model (1,000 trees, maximum 20 nodes) was trained on the complete dataset using these labels to learn optimal feature weights. The model integrated normalized features—including, z-scored metrics, scaled maximum differences, and binary specificity flags—into a cell type specificity score (CSS) for all genes. Genes expressed in >3%, >2%, or >1% of cells in a cell type population (for Levels-3, -2, and -1, respectively) were used for statistical testing. Statistical significance was assessed by fitting a beta distribution to background scores from genes not meeting the specificity criteria and computing FDR-adjusted p-values. Genes were classified as cell type-specific or globally expressed at adjusted p-value < 0.05.

*Origin-of-change analysis.* To distinguish changes in cell abundance from cell-intrinsic expression changes occurring with aging, we used TMS spleen data from ages 1, 3, 18, 21, 24, and 30 months. Using linear regression, we tested age effects on three features per gene per cell type: total expression (sum of normalized expression across all cells), number of cells expressing the gene, and fraction of cells expressing the gene. Data were rank-z normalized per gene and linear models were fitted as Model 9: Y ~ Age, where Y represents the rank-z normalized feature and Age is continuous in months. Age coefficients, scaled coefficients, and p-values were extracted per feature. Correction for multiple testing was performed per cell type using the Benjamini-Hochberg procedure. Gene features were classified as significantly changing at adjusted p-value < 0.1.

*Benchmarking.* Celestial marker performance was benchmarked against Seurat-derived markers[78] at both cell-level and bulk-level resolution. Cell-level accuracy was assessed by computing mean log-normalized marker expression per cell and calculating the area under the precision-recall curve (AUPRC) against binary cell type labels across all TMS droplet spleen cells. Bulk-level accuracy was assessed by constructing per-cell type pseudobulks per sample—the sum of raw counts within each cell type, normalized by total counts per sample—resulting in sample-level CPM values comparable to bulk RNA-seq measurements. AUPRC was determined by testing whether each cell type's mean marker CPM ranked above all other cell types across samples. Two Seurat settings were used for comparison: a stringent setting (min.pct = 0.03, logfc.threshold = 4), designed to match Celestial's minimum fraction threshold and select only highly specific markers; and a standard setting (min.pct = 0.25, logfc.threshold = 1), representing standard single-cell marker calling.

*Cross-species analyses of splenic gene-to-cell-type assignments.* Celestial was applied to human scRNA-seq studies to assess the spleen markers across datasets and between species. The standard settings for Celestial were used [19,68–70].
